# Structure-based design of a highly stable, covalently-linked SARS-CoV-2 spike trimer with improved structural properties and immunogenicity

**DOI:** 10.1101/2021.05.06.441046

**Authors:** Eduardo Olmedillas, Colin J. Mann, Weiwei Peng, Ying-Ting Wang, Ruben Diaz Avalos, Dan Bedinger, Kristen Valentine, Norazizah Shafee, Sharon L. Schendel, Meng Yuan, Guojun Lang, Romain Rouet, Daniel Christ, Weidong Jiang, Ian A. Wilson, Tim Germann, Sujan Shresta, Joost Snijder, Erica Ollmann Saphire

## Abstract

The continued threat of SARS-CoV-2 to global health necessitates development of improved research tools and vaccines. We present an improved SARS-CoV-2 spike ectodomain, “VFLIP”, bearing five proline substitutions, a flexible cleavage site linker, and an inter-protomer disulfide bond. VFLIP displays significantly improved stability, high-yield production and retains its trimeric state without exogenous trimerization motifs. High-resolution cryo-EM and glycan profiling reveal that the VFLIP quaternary structure and glycosylation mimic the native spike on the viral surface. Further, VFLIP has enhanced affinity and binding kinetics relative to other stabilized spike proteins for antibodies in the Coronavirus Immunotherapeutic Consortium (CoVIC), and mice immunized with VFLIP exhibit potent neutralizing antibody responses against wild-type and B.1.351 live SARS-CoV-2. Taken together, VFLIP represents an improved tool for diagnostics, structural biology, antibody discovery, and vaccine design.

## INTRODUCTION

The likely persistence of SARS-CoV-2 necessitates development of improved research tools, diagnostic reagents, and next-generation vaccine candidates. Vaccine approaches in progress involve inactivated virus, viral vectors, protein subunits, and liposome-encapsulated mRNA (Krammer 2020). Despite their different modes of delivery, all of these platforms use the SARS-CoV-2 surface glycoprotein spike as the primary target. Spike is a metastable, membrane-anchored, class I fusion protein that exists as a trimer on the viral surface. A multi-basic furin cleavage site (residues 682-685) delineates the boundary between the spike S1 (receptor-binding) and S2 (membrane fusion) domain subunits. A second cleavage site, S2’, lies downstream of the furin site and is cleaved by TMPRSS2 on the target cell surface or cathepsin L following endocytic uptake. Cleavage at both S1/S2 and S2’ facilitates liberation of the fusion peptide and triggering of structural rearrangements that promote the pre-fusion to post-fusion transition and drive virus and host membrane fusion (Shang, Wan, Luo, et al. 2020; Sacco et al. 2020). The inherent instability of the prefusion state complicates use of spike and spike ectodomains in vaccines, diagnostics, and research applications as presentation of the native quaternary structure is critical for eliciting and detecting potently neutralizing conformation-dependent antibodies (Liu et al. 2020; Cai et al. 2020). Substantial effort has been devoted to developing versions of the spike glycoprotein that retain its immunologically relevant, native-like prefusion state.

On the virion surface, ~97% of the spikes exist in the metastable pre-fusion conformation (Ke et al. 2020). While in this pre-fusion state, the S1 domains of the spike exhibit dynamic motion. The receptor-binding domains (RBDs) that lie at the apex of the spike are begin predominantly in a “down” conformation with the receptor-binding motif (RBM) buried by RBD-RBD interfaces that together form a trefoil structure (Lu et al. 2020; Yao et al. 2020). Transiently, or upon interaction with the angiotensin-converting enzyme 2 (ACE2) receptor on the target cell surface, the RBDs lift up sequentially and the corresponding N-terminal domains (NTD) undergo rotational movements. Transition of the RBD to the “up” state reduces the contact surface area between adjacent RBDs and with the NTD, eventually leading to dissociation of S1 from the S2 subunit and promoting S2 transition to the post-fusion state (Lu et al. 2020). Engineered spikes used for immunogens and research tools should not only preserve the prefusion trimeric assembly, but also the native positioning of RBDs and RBD motion. In addition, accurate reflection of glycans presented on the native virion is key for eliciting and detecting antibodies induced by infection or vaccination (He et al. 2018; Cao et al. 2017; Sun et al. 2013; Grant et al. 2020).

Vaccines currently deployed in the United States use a derivative of the prototypical, first-generation “S-2P” spike design (Pallesen et al. 2017), which contains two proline substitutions at positions 986 and 987 (Polack et al. 2020; Bos et al. 2020; Corbett et al. 2020; Wrapp et al. 2020). These vaccines have shown high efficacy in the short term, but the rapid timeframe for development has afforded few opportunities for antigen optimization. Recent work by Hsieh *et al*. illustrated that the S-2P spike exhibits relatively low yield and unfavorable purity (Hsieh et al. 2020). The poor yield may impact cost and manufacturability of vaccine candidates, and limit expression levels in vaccinated individuals, which in turn could necessitate higher doses and potentially increased reactogenicity. Moreover, several studies reported that S-2P protein preparations exhibit sensitivity to cold-temperature storage (Edwards et al. 2020; Xiong et al. 2020). Edwards et al. used negative-stain electron microscopy (NSEM) to demonstrate a 95% loss of well-formed S-2P spike trimers after 5-7 days of storage at 4°C. Exposure to 4°C temperatures also resulted in lower thermostability and altered binding to monoclonal antibody (mAb) CR3022, suggesting perturbed structure and antigenicity.

A second-generation spike construct, termed “HexaPro”, contains four additional prolines at positions 817, 892, 899 and 942. HexaPro expresses to levels nearly 10-fold higher than those for wild-type spike or S-2P, has a 5 °C higher melting temperature (Tm) (Hsieh et al. 2020), and displays improved stability relative to S-2P under low-temperature storage and multiple freeze-thaw cycles (Edwards et al. 2020). Importantly, binding assays and cryoEM indicated that HexaPro better retains the native prefusion quaternary structure compared to S-2P, despite still exhibiting minor reductions in thermostability and mAb binding following incubation at 4 °C.

Both S-2P and HexaPro, however, are prone to antibody-and ACE2-mediated triggering of conformational change to the post-fusion state (Huo et al. 2020; Ge et al. 2021; Xiong et al. 2020). This triggering complicates structural analysis of mAb-spike and ACE2-spike complexes and may affect immunogenicity upon vaccination. Several spike constructs such as SR/X2 prevent this fusogenic activity with introduction of an inter-protomer disulfide bond, linking the RBD and the S2 subunits to “lock” the RBDs in the “down” conformation (Xiong et al. 2020; Henderson et al. 2020). Although these “locked-down” spike proteins maintain the trimeric state, the location of the inter-protomer disulfide bond prevents the natural hinge motion of the RBD and ablates binding to ACE2 and “RBD-up” antibodies, which are among the most potent neutralizers (Rogers et al. 2020; Liu et al. 2020; Huo et al. 2020; Brouwer et al. 2020). Furthermore, cryo-EM structures of a locked-down spike show that the RBDs are rotated 2Å closer to the three-fold axis relative to wildtype (Xiong et al. 2020). These quaternary structure perturbations, together with locking of the RBD into an “all-down” state, could prevent elicitation and detection of protective antibodies against neutralizing epitopes that are only accessible in the “up” or mixed up/down conformation (Rogers et al. 2020; Huo et al. 2020; Liu et al. 2020; Brouwer et al. 2020). Even antibodies that target the “all-down” RBD conformation could be affected, particularly those that bridge two RBDs, such as the potent neutralizing mAbs S2M11, Nb6, and C144 (Schoof et al. 2020; Tortorici et al. 2020; Robbiani et al. 2020). Thus, a spike immunogen that preserves the natural RBD positioning and conformational dynamics is essential for maintaining the native antigenic landscape.

A central goal for SARS-CoV-2 vaccines is to reduce incidence of symptomatic disease through generation of enduring protective immunity. However, the recent emergence of SARS-CoV-2 variants of concern (VOC) poses a risk to first-generation vaccine efficacy and durability of both infection-and vaccine-induced humoral immunity. Lineage B.1.351 (informally known as the South African variant) is particularly concerning due to substitutions that confer increased transmissibility and reduced sensitivity to neutralization by heterotypic convalescent and vaccine-induced sera. Development of structurally designed vaccine candidates with improved immunogenicity and breadth of coverage is critical for controlling emergent VOC.

To address these issues associated with current spike constructs and emergence of VOC, we performed iterative design of spike proteins containing different proline substitutions, cleavage site linkers, and interprotomer disulfide bonds. We describe the production of “VFLIP” (five (V) prolines, Flexibly-Linked, Inter-Protomer disulfide) spikes that remain trimeric without exogenous trimerization motifs, and which have enhanced thermostability relative to earlier spike constructs. Surface plasmon resonance (SPR) and cryo-EM analysis confirm the native-like antigenicity of VFLIP and its improved utility for structural biology applications. Moreover, mice immunized with the VFLIP spike elicited significantly more potent neutralizing antibody responses against live SARS-CoV-2 D614G and B.1.351 compared to those immunized with S-2P. Taken together, our data indicate that VFLIP is a thermostable, covalently-linked, native-like spike trimer that represents a promising next-generation research reagent, diagnostic tool, and vaccine candidate.

## RESULTS

### Iterative design of the VFLIP spike immunogen

In the SARS-CoV-2 spike, the S1/S2 furin cleavage site is located in a flexible external loop (residues 675-691) that can enhance triggering of the post-fusion conformation (Shang, Wan, Liu, et al. 2020). However, the sequence of this loop varies among coronavirus species and is absent in the closely related SARS-CoV and RaTG13 (Walls, Park, et al. 2020)”. Considering this lack of sequence conservation and role in trimer metastability, we hypothesized that replacing 15-aa of the loop (residues 676-690) with a shorter, rigid (Gly-Pro) or flexible (Gly-Gly-Gly-Ser) linker may improve stability and yield without disturbing the quaternary structure or native antigenicity (Figure 1A and Supp. Table 2). Five constructs bearing linkers of differing rigidity and length were evaluated. Constructs 1 and 2 have flexible Gly-Gly-Gly-Ser linkers (termed FL1 and FL2 for Flexible Linker) and constructs 3-4 have rigid Gly-Pro linkers (RL3-RL4 for Rigid Linker) (Supp. Table 1). Expression of SARS S-2P_FL1 or S-2P_FL2 resulted in a 2-4 fold greater yield of trimeric spike over S-2P in both HEK293F and ExpiCHO cells (Supp. Table. 1).

**Figure 1.**
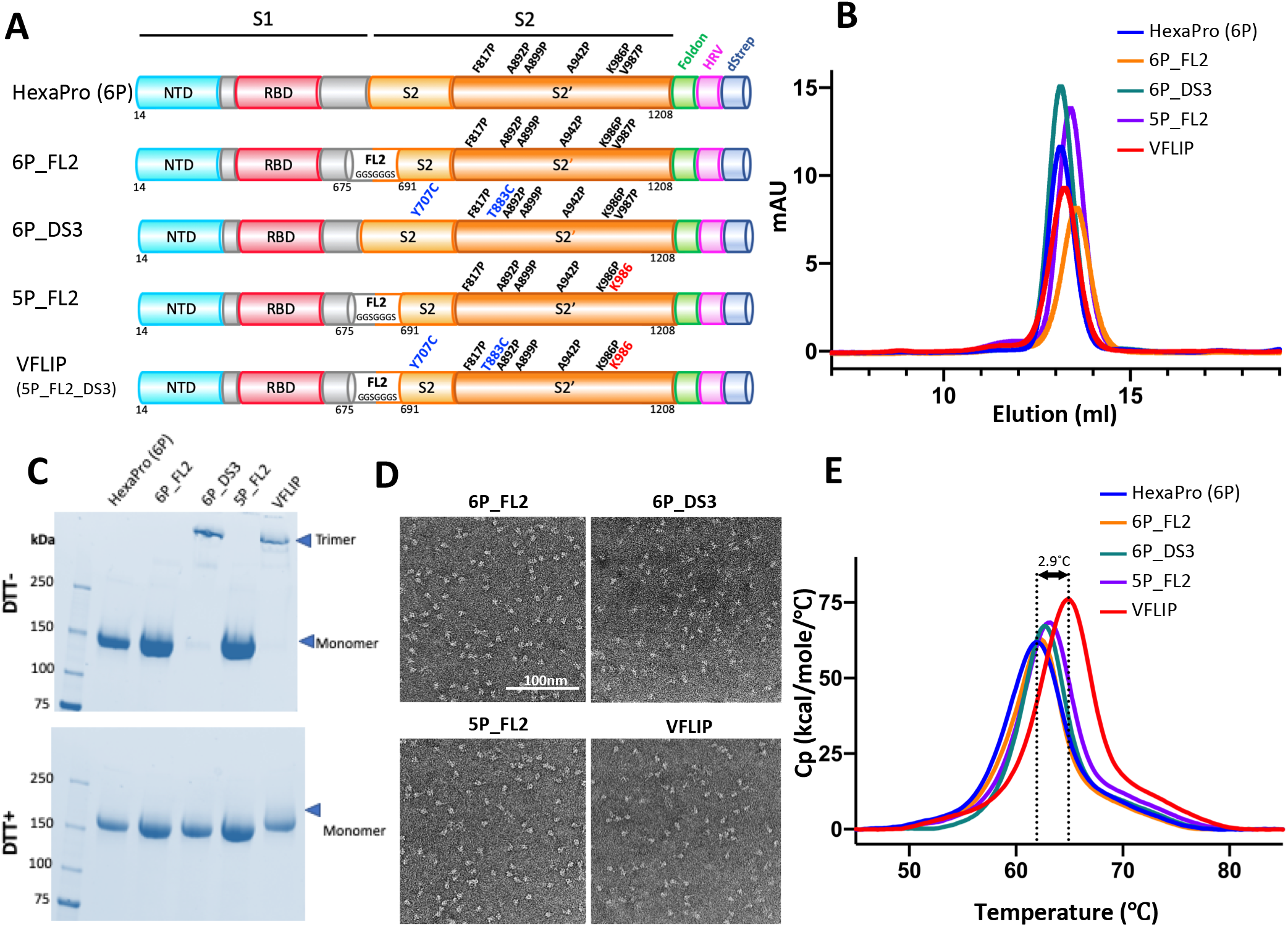
Expression and characterization of a covalently-linked thermostable spike variant. **(A)** Schematic representation of VFLIP and intermediate constructs indicating location of engineered mutations and structural features. Cysteine point mutations to introduce a disulfide bond are highlighted in blue text, while restoration of lysine at position 986 (K986) is highlighted in red text. The flexible linker (FL) inserted in some constructs is shown in white. **(B)** Analytical size-exclusion chromatography comparing spike purity post-purification. **(C)** Non-reducing/Reducing SDS-PAGE of constructs shown in (A). **(D)** Representative negative-stain electron micrographs of VFLIP and intermediate constructs showing homogenous well-formed trimer populations. **(E)** Differential Scanning Calorimetry (DSC) analysis of spike thermostability. Arrows indicate increase in Tm for VFLIP relative to HexaPro.

We next explored the relative impact of these linkers in the context of the six-proline (6P)-bearing spike, with the six prolines representing those in HexaPro (Hsieh et al. 2020). The 6P_RL constructs bearing rigid linkers had slightly lower expression levels relative to HexaPro. In contrast, the flexibly-linked 6P_FL1 and 6P_FL2 constructs had approximately equivalent or increased yields compared to HexaPro, with 6P_FL2 averaging 15.6 mg/L in HEK293F and 184 mg/L in ExpiCHO cells. Dynamic scanning calorimetry (DSC) analysis showed a ∼1 °C increase in melting temperature (Tm) for all tested linkers (flexible or rigid) relative to the parental HexaPro (Figure 1E). The FL2 linker, which had the best performance in terms of yield and stability, was selected for further use.

Both S-2P and HexaPro have a proline substitution at residue 986 (K986P). However, multiple studies have suggested a role for K986 in mediating the RBD conformational state and S2 stability via electrostatic interactions with D427 (Xiong et al. 2020; Cai et al. 2020; Gobeil et al. 2021) and/or D748 (Juraszek et al. 2021), respectively. Juraszek et al. also showed that spikes bearing the K986P mutation have a substantially larger fraction of “RBD-up” spikes and reduced thermostability. We therefore modified the six-proline-containing FL2 spike (6P_FL2) by reverting the K986P mutation back to wild-type K986 to generate the “5P_FL2” construct. When transfected in HEK293F cells, 5P_FL2 expresses an average of 40% higher than HexaPro and 23% higher in ExpiCHO cells. DSC indicates that 5P_FL2 indeed has greater thermostability than 6P_FL2 (Figure 1E), consistent with Juraszek et al. (2021). The 5P_FL2 construct allows production of highly pure, well-formed trimers as confirmed by size-exclusion-chromatography (SEC) (Figure 1B) and negative-stain electron microscopy (NSEM) (Figure 1D).

Introduction of disulfide bonds can substantially increase the stability of viral glycoproteins, as has been demonstrated for HIV, LASV and RSV (Stewart-Jones et al. 2015; Hastie et al. 2017; McLellan et al. 2013). We identified and tested a set of eight inter-protomeric disulfide bonds (DS1-DS8) for their ability to improve stability of the trimer. After an initial screening in the S-2P background, the best disulfide bond candidates were transferred to the 6P background for further analysis. Among the disulfide-linked spikes, the 6P_DS3 construct (Y707C/T883C) displayed the best yield and quality (purity and homogeneity of oligomerization state) following purification (Figure 1C, Supp. Table S1). When analyzed by non-reducing SDS-PAGE, the 6P_DS3 construct retains its covalently-linked trimeric state (>250 kDa) without detectable low molecular weight species, indicating efficient disulfide bond formation. Upon reduction with DTT, 6P_DS3 runs at the monomeric molecular weight of the non-disulfide linked constructs (∼180 kDa). SEC and NSEM confirmed that 6P_DS3 expresses as a highly homogeneous population of well-formed pre-fusion spike trimers. Although addition of the DS3 disulfide bond reduces total yield ∼12% relative to 6P alone, it does increase thermostability slightly (0.5 °C increase in melting temperature) as measured by DSC, but more importantly, robustly maintains its trimeric state (Figure 1E).

Given the improvements in yield and stability afforded by the 5P background, the flexible FL2 linker, and the DS3 disulfide bond individually, we then generated 5P_FL2_DS3, which combines the three modifications. This construct is hereafter termed VFLIP spike for 5 (V) proline, Flexibly-Linked, Inter-Protomer bonded spike. The VFLIP spike exhibits the best thermostability of the constructs tested here, with a Tm that is 2.9 °C higher than that for HexaPro (Figure 1E). VFLIP is efficiently expressed in both HEK293F and ExpiCHO cells, with an average yield of 7.1 mg/L and 157 mg/L, respectively. Hence, the VFLIP spike was deemed a promising antigen suitable for in-depth characterization.

### Structural characterization of the engineered VFLIP and VFLIP_D614G spikes

We analyzed VFLIP by high-resolution cryo-electron microscopy (cryo-EM) to understand the molecular mechanisms underlying its increased stability. We also obtained a high-resolution cryo-EM structure for a VFLIP variant bearing the now globally fixed D614G mutation (Korber et al. 2020). From a single VFLIP dataset, we identified a population of 166,085 particles, to produce a density map having an overall resolution of 3.0 Å with no applied C3 symmetry (Figs. 2A and Supp. Figure S1). For VFLIP_D614G, a population of 213,852 particles yielded a 2.8Å resolution structure (Figure S2). Importantly, the density maps confirm the successful formation of the DS3 disulfide bond (Y707C/T883C) and the K986/E748 salt-bridge interaction (Figure 2B and Supp. Figure S3).

**Figure 2.**
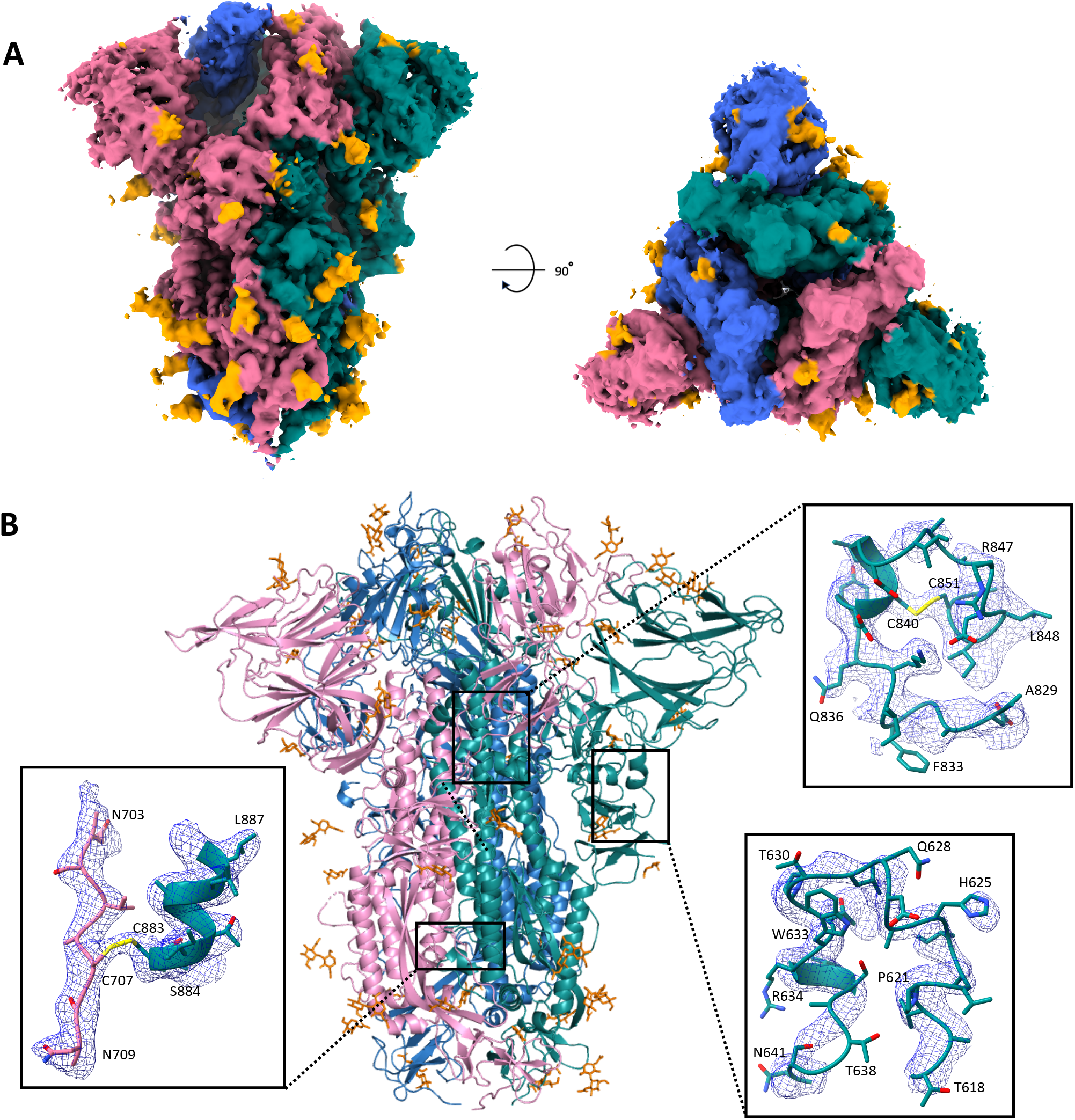
High resolution Cryo-EM structure of VFLIP. **(A)** Side (left) and top (right) views of the electron density map for VFLIP Wuhan variant (D614). Individual protomer surfaces are shown in pink, green or blue, and N-linked glycan densities are shown in orange. **(B)** Ribbon structure of VFLIP using the same color scheme as in (A). The expanded view in the lower left panel shows the density and corresponding model of the Y707C/T883C inter-protomer disulfide-bond in yellow. Expanded view in the top right panel shows density and modeling of the 829-852 loop, including the C840/C851 disulfide bond (yellow). Density and a model for the 618-641 loop, often not visible in previous spike ectodomain structures, is shown in the expanded view in the panel on the lower right.

Extensive classification and data processing reveal >90% predominantly closed trimers, with all RBDs “down” for both VFLIP and VFLIP_D614G. Further sub-classification revealed an overall architecture that is similar to other “closed” spikes in the Protein Data Bank (PDB). The improved order of this construct allows visualization of portions of the SARS-CoV-2 spike that are rarely visible in other cryoEM structures. For example, the peptide backbone of the region extending from residues 618-641, which fits/wedges between RBD and NTD, can now be traced (Figure 2B). The fusion peptide proximal region (FPPR) extending from residues 829-851 that contains the C840-C851 disulfide bond, is also now well-ordered in VFLIP (Figure 2B). This loop packs against the S2 helical core and nestles beside the 630 loop and underneath CTD2 of the adjacent protomer to provide stability through pi-stacking interactions (Benton et al. 2020).

### Native-like glycosylation of VFLIP expressed in HEK293F and ExpiCHO cells

A protein used as an immunogen or as a research tool to detect antibodies should present the native glycosylation profile (Grant et al. 2020; Sun et al. 2013; Raska et al. 2010; Watanabe et al. 2020; He et al. 2018). In SARS-CoV-2 spike, glycans play a major role in RBD positioning and trimer stability (Casalino et al. 2020; Sztain et al. 2021). Thus, recapitulating native glycoforms is likely important to preserve antigenicity. We used liquid chromatography coupled with electron transfer/high energy collision dissociation tandem mass spectrometry (LC-MS/MS) to determine the glycosylation profiles of VFLIP produced in HEK293T and ExpiCHO cell lines, and compared these profiles to previously published data for S-2P and virion-associated spikes (Yao et al. 2020; Watanabe et al. 2020). We detected N-linked glycopeptides at 20 of 22 predicted glycosylation sequons (with N17 and N1158 not covered in our experiments), and measured the associated diversity in glycan processing (Figure 3A). A full overview of the site-specific individual glycoforms is presented in Supplementary Figure S4. Similar to native virions and S-2P, the glycosylation pattern in both 5P and VFLIP spikes is dominated by complex glycosylation, with selected sites enriched in underprocessed, high-mannose glycans. The pattern of glycan processing of VFLIP produced in HEK293F or ExpiCHO cells was highly similar. Glycans of the S1 subunit, including those located within the RBD (N331 and N343) and those modulating its up/down conformation (N165 and N234), were similarly processed between VFLIP spikes or 5P spikes. Likewise, glycans at the base of the S2 subunit (N1074, N1098, N1134, N1173 and N1194) were similar in 5P and VFLIP spikes. However, N-glycan patterns for 5P and VFLIP differed near the middle of the spike, particularly for glycans N603, N616, N709, and N801, which were largely underprocessed in 5P spikes (Figs. 3A-C). Previous glycan analysis of the S-2P spike revealed similar underprocessing at these glycan positions compared to the native virion (Frese et al. 2012, 2013), where these glycan sites were found to be predominantly complex-type and more similar to VFLIP produced from both HEK293F and ExpiCHO (Figure 3C). Taken together, these data indicate that VFLIP spikes more accurately reflect glycosylation patterns of the native spike, and that VFLIP spikes produced in HEK293F and ExpiCHO have nearly identical glycan composition.

**Figure 3.**
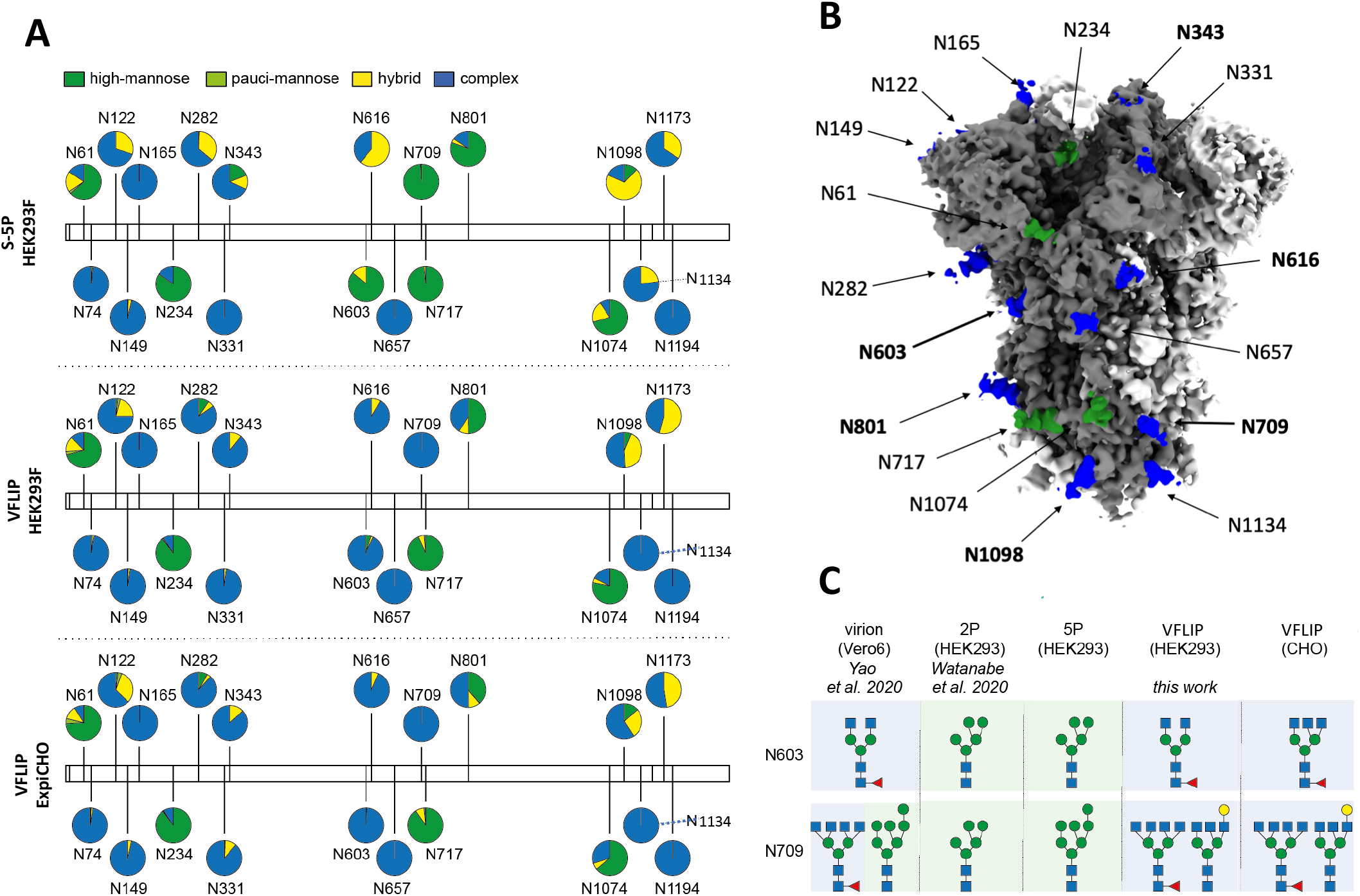
Site-specific glycan analysis of VFLIP and 5P spikes. **(A)** Schematic illustration of the quantitative mass spectrometric analysis of the glycan population present on VFLIP produced in HEK293F and ExpiCHO compared to a spike construct containing five proline substitutions (5P) produced in HEK293F. Pie charts show the glycoform distribution detected at each glycan site **(B)** Electron density map of trimeric VFLIP with N-linked glycans colored according to complexity as in (A). Bold text indicates glycans that show distinct differences in processing between VFLIP and 5P. **(C)** Similar processing at N603 and N709 in VFLIP and virion-associated spikes that differs from processing seen in previously described proline-stabilized constructs. Blue square, NAG. Green circle, mannose. Red triangle, fucose. Yellow circle, galactose.

### Epitope mapping and binding kinetics of over 100 mAbs

To verify whether introduced modifications could affect functional epitopes, we probed VFLIP antigenicity by analyzing recognition of 136 monoclonal antibodies (mAbs) from the Coronavirus Immunotherapeutic Consortium (CoVIC). HexaPro, which shows a high proportion of one or more RBDs in the up state, and 5P_FL5_DS2 (referred to here as DS2), a locked-down spike, were used for comparison, along with soluble RBD. High-throughput surface plasmon resonance (SPR) based assays revealed distinct IgG affinities for each analyzed antigen (Figure 4A). Both the association rate (*k*a) and apparent dissociation rate (*k*d) of the mAbs for VFLIP were slightly slower than that for HexaPro, but the resulting sub-nanomolar affinities were similar between VFLIP and HexaPro for most mAbs. The slower association rate observed for some VFLIP-antibody interactions may be due to the predominantly “all-down” initial conformation and the additional stability provided by the well-ordered 630 loop and FPPR that likely slow RBD state transition. The slower dissociation constant, however, restores overall affinity and may facilitate isolation of antibody:spike complexes for structural analyses.

**Figure 4.**
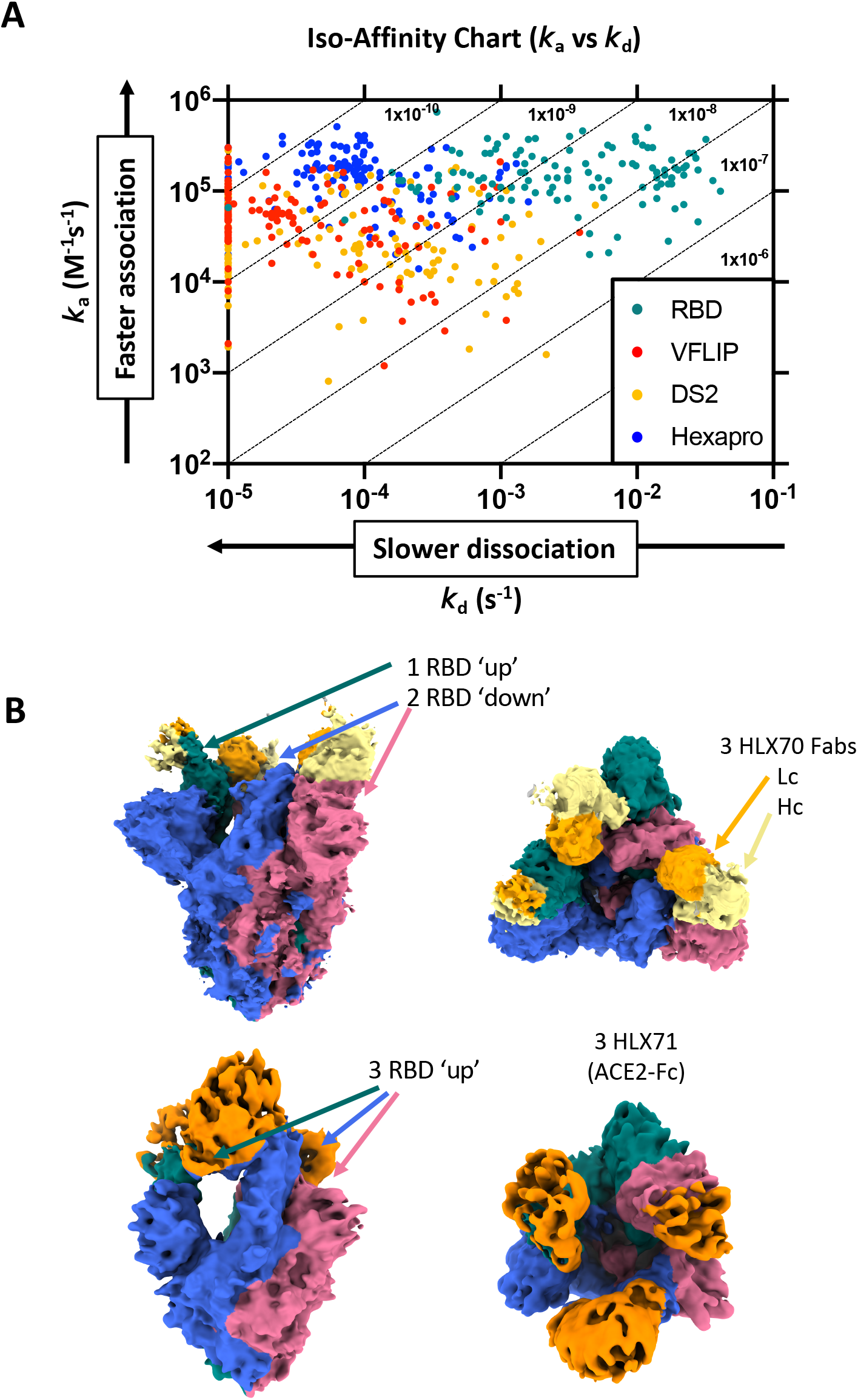
Binding kinetic analysis with over 100 CoVIC monoclonal antibodies. **(A)** IgG affinities for 136 mAbs from the CoVIC consortium for soluble RBD (green), VFLIP (red), 5P.USEO5.DS2 (yellow), and HexaPro (blue). Y-axis shows the association rate (*k*_a_) and the arrow indicates increasing association rate. The X-axis shows the dissociation rate (*k*_d_) and the arrow indicates decreasing dissociation rates. Diagonal dashed lines delineate calculated molar affinities (*K*_D_) given in log-scale. **(B)** Cryo-EM density maps of VFLIP_D614G complexed with a Fab from HLX70, a humanized pan-coronavirus mAb (Top), and HLX71, an ACE2-Fc fusion protein (bottom). Spike protomers are colored in pink, blue and green. HLX70 Fab is bound in a unique “1-up, 2-down” conformation. Fab heavy and light chains are shown in yellow and orange, respectively. ACE2-Fc is bound to VFLIP.D614G (orange) with a fully occupied stoichiometry of three molecules per spike.

In contrast, around 30% of the mAbs tested had at least 10-fold lower affinity for the locked-down DS2 spike relative toVFLIP. As expected, soluble monomeric RBD exhibited the highest association rate for most antibodies, but also had a much higher dissociation rate, and no potential for avidity as seen with the trimers, and thus bound to mAbs with 10-100 fold lower apparent affinity. Overall, our data suggests that VFLIP maintains native-like antigenicity and binding to mAbs of potential clinical interest, further supporting its utility as a candidate spike antigen.

Considering that unbound VFLIP is mostly in a closed conformation, yet still binds to RBD-up antibodies, we sought to determine the structure of VFLIP in complex with RBD-binding mAbs. We obtained high-resolution cryo-EM structures of VFLIP in the “up” conformation by complexing with a Fab from the therapeutic candidates HLX70 (mAb) and HLX71 (an ACE2-Fc fusion protein) (Figure 4B and Suppl. Figs. S5 and S6). Both candidates are currently in clinical trials (NCT04561076 and NCT04583228, respectively) (“Evaluate Safety and Pharmacokinetics of HLX70 in Healthy Adult Volunteers” n.d.)(“Evaluate the Safety, Tolerability, Pharmacodynamics, Pharmacokinetics, and Immunogenicity of HLX71 (Recombinant Human Angiotensin-Converting Enzyme 2-Fc Fusion Protein for COVID-19) in Healthy Adult Subjects” n.d.). Single-particle reconstruction of the VFLIP-HLX70 Fab complex produced a 3.6 Å structure that revealed a unique conformation and a fully occupied 3:1 ratio for Fab:spike, with one RBD “up” and two RBDs “down” (Figure 4B). For the VFLIP-ACE2-Fc complex, the major population displayed three RBDs in the up-state with full occupancy by three ACE2 molecules. This structure was resolved to 4.6 Å. The stoichiometry of one VFLIP bound to three copies of ACE2 is consistent with previous ACE2-bound structures, and demonstrates that VFLIP modifications do not abrogate RBD re-positioning or binding to RBD-up antibodies.

### VFLIPΔFoldon remains a stable trimer under diverse biochemical conditions

Most soluble versions of the SARS-CoV-2 spike have a T4 phage fibritin trimerization motif (termed Foldon, PDB ID: 1RFO) at the C-terminus of the ectodomain (Güthe et al. 2004; Wrapp et al. 2020; Henderson et al. 2020; Xiong et al. 2020; Hsieh et al. 2020). Foldon is, however, immunogenic and may influence the anti-spike immune response (Sliepen et al. 2015; Lainšček et al. 2020). Thus, we designed a VFLIP construct containing an HRV-3C site between the C-terminus of the spike and the Foldon domain that allowed its selective removal following purification to produce a protein termed VFLIPΔFoldon (Figure 5A).

**Figure 5.**
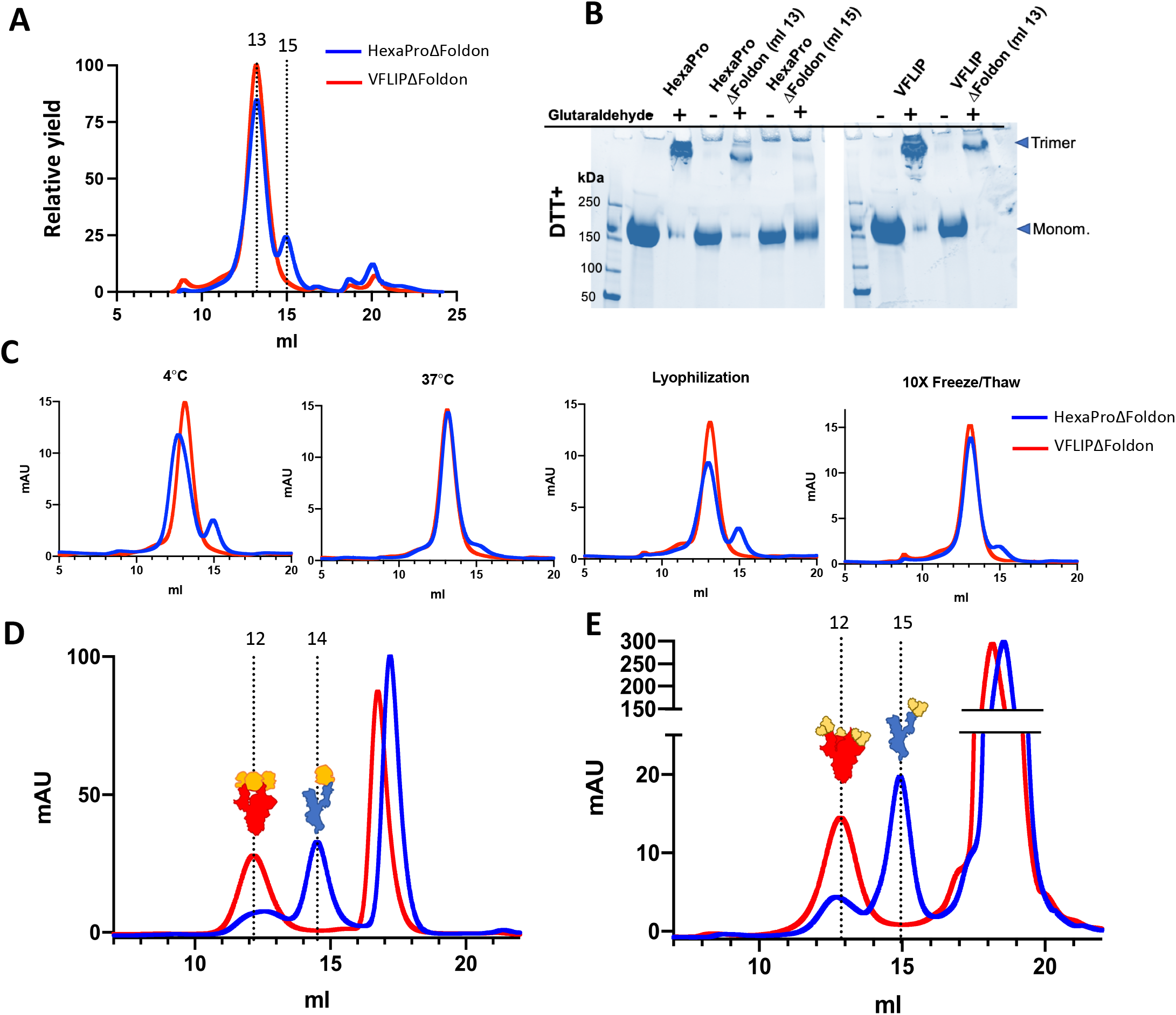
Biophysical stability of VFLIPΔFoldon and resistance to receptor-induced dissociation. **(A)** SEC trace of HexaPro and VFLIP spikes following removal of the Foldon domain (ΔFoldon). **(B)** Oligomeric state of peak fractions evaluated by reducing SDS-PAGE with and without prior glutaraldehyde crosslinking. **(C)** SEC of HexaProΔFoldon and VFLIPΔFoldon following incubation for five days at 4°C or 37 °C, lyophilization/reconstitution, or ten freeze/thaw cycles. VFLIPΔFoldon retains trimeric assembly (13 mL), whereas a portion of HexaΔFoldon dissociates to monomers (15 mL). **(D, E)** SEC trace of VFLIPΔFoldon (red) and HexaProΔFoldon (blue) complexed with soluble human ACE2 (D) and B6 Fab (E). VFLIPΔFoldon remains trimeric after binding to ACE2 or B6 Fab, but a majority of HexaProΔFoldon trimers dissociate to monomers.

To determine the impact of removing the Foldon domain on spike stability, we subjected VFLIPΔFoldon to biochemical stress tests. Initial SEC analysis of purified VFLIPΔFoldon showed a narrow single peak centered at 13 ml with no detectable aggregation or lower molecular weight species (Figure 5A). Under the same proteolytic treatment, HexaProΔFoldon consistently eluted as a double peak (13 ml and 15 ml), with 15% of the total mass corresponding to the lower molecular weight species (Figure 5A).

VFLIPΔFoldon also remains stable after prolonged (> 1 month) storage at room temperature, 4 °C and 37 °C (conditions that cause a portion of HexaPro trimers to dissociate into species having a lower molecular weight consistent with that for monomers), as well as lyophilization and multiple freeze/thaw cycles (Figure 5C). Thus, in addition to improved stability under standard purification conditions, VFLIP exhibits resistance to adverse conditions, even without an exogenous trimerization domain.

### VFLIP enables structural characterization of receptor-mimicking antibodies and ACE2

ACE2 and many RBD-targeted antibodies exhibit fusogenic activity and trigger transition of the spike to the post-fusion structure upon binding (Huo et al. 2020). Various methods have been implemented to circumvent this issue, such as use of cross-linking agents (Xu et al. 2021) or minute-scale incubation prior to freezing grids (Benton et al. 2020). However, such approaches often produce samples having poor or no occupancy that complicates structural determination (Benton et al. 2020; T. Zhou et al. 2020). Some studies successfully identified fully occupied HexaPro:ACE2 complexes, but these particles were rare and represented as little as 3% of the total particle population (Benton et al. 2020).

To address this issue, we generated complexes of soluble human ACE2 (shACE2) with HexaProΔFoldon or VFLIPΔFoldon and purified them by SEC (Figure 5D). For HexaProΔFoldon, only ~15% of the total mass corresponded to trimeric spike bound to ACE2, and the remainder eluted as monomeric S bound to ACE2 at 14 ml. In contrast, VFLIP:ACE2 complexes eluted as a single peak at 12 ml, indicating consistent occupancy and lack of trimer dissociation. This result corresponds to our high-resolution cryo-EM structure showing three ACE2 molecules bound to each spike (Figure 4B). Multiple studies have noted that some antibodies against the RBD “split” spike into monomers (Walls, Xiong, et al. 2020; Koenig et al. 2021). The mAb B6 is one such spike-splitting antibody. When the Fab of B6 is complexed with HexaProΔFoldon, ∼90% of the HexaPro spike separates to monomers. In contrast, complexes of B6 Fab with VFLIPΔFoldon retain the trimeric spike assembly, and exist in complete occupancy, one spike trimer to three Fab fragments (Figure 5E).

### VFLIP elicits potently neutralizing responses in immunized mice

We hypothesized that the additional stability afforded by VFLIP may improve elicitation of neutralizing antibodies and that removing Foldon (VFLIPΔFoldon) may avoid deleterious responses to this exogenous trimerization domain. To assess the immunogenicity of VFLIP, we immunized BALB/c mice with four versions of spike: (1) Parental S-2P, (2) HexaPro, (3) VFLIP, and (4) VFLIPΔFoldon. Ten mice per group were immunized intramuscularly (i.m.) with 25 μg purified spike adjuvanted with CpG + alum and boosted with the same four weeks later. An interim blood draw was performed two weeks after the prime, and a final blood draw was taken two weeks after the boost. Overall, mice in all five groups mounted robust antibody responses as evidenced by total anti-spike antibody titers (Figure 6B). Sera from immunized mice were used to examine activity in neutralization assays using rVSV-pseudotyped with SARS-CoV-2 spike bearing the D614G mutation, as well as authentic SARS-CoV-2 bearing the D614G substitution and authentic SARS-CoV-2 virus of the B.1.351 (South African) lineage, which is a current variant of concern. Pseudovirus neutralization titers for VFLIP-immunized sera were somewhat higher than HexaPro and achieved 50% neutralization at dilutions over 1:100,000. Assays using authentic D614G and B.1.351 showed that VFLIP-induced sera had a higher neutralizing potency compared to S-2P, with 50% neutralization at dilutions of 1:30,000 and 1:13,000, respectively (Fig, 6 D,E). Pseudovirus neutralization of VFLIP-ΔFoldon (with trimerization domain removed) was equivalent to that for Foldon-containing HexaPro, indicating that immunogenicity is maintained without an exogenous trimerization motif (Figure 6C).

**Figure 6.**
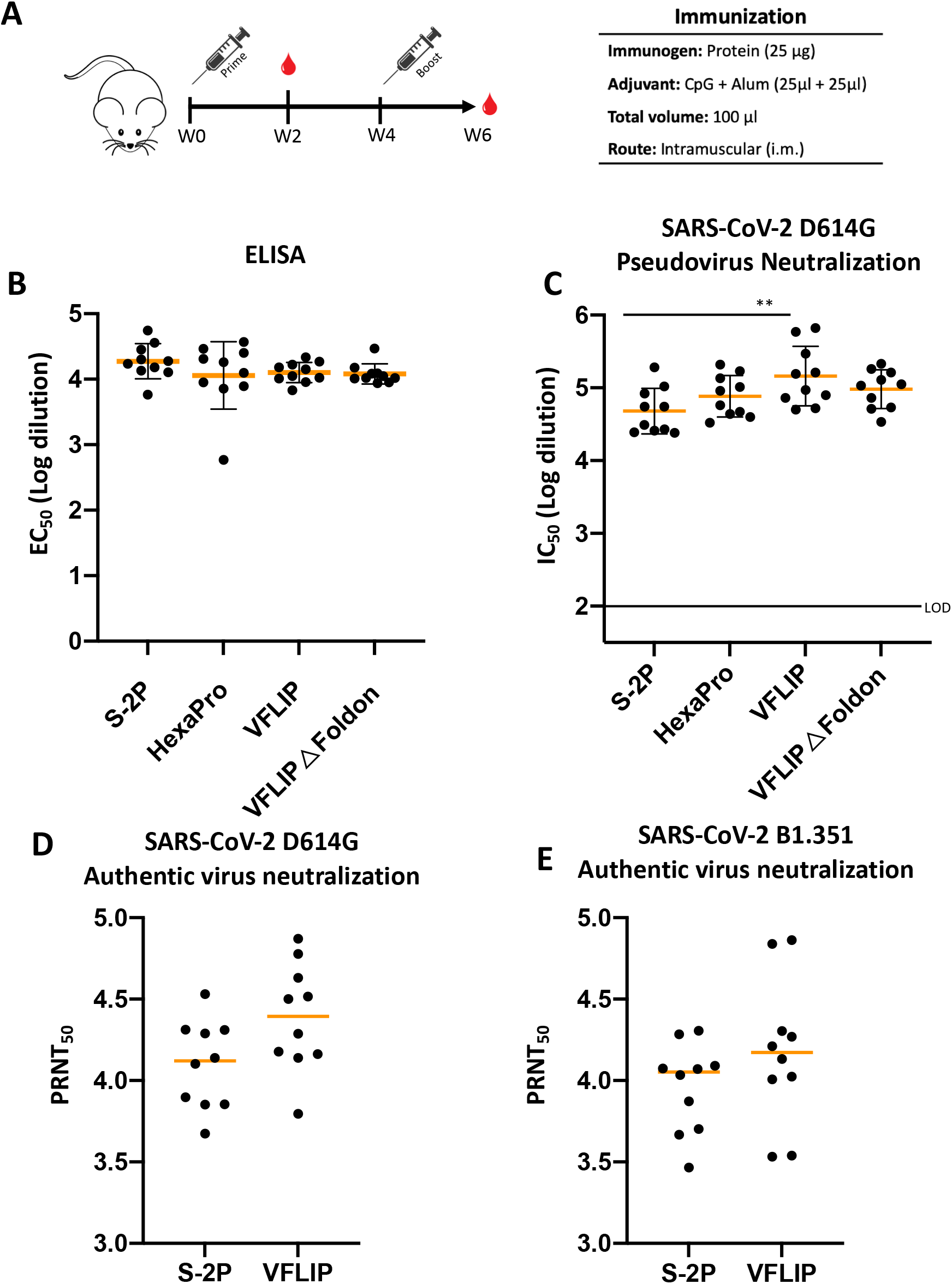
Immunogenicity of VFLIP spikes in BALB/c mice. **(A)** Schematic representation of the mouse immunization protocol. Mice were immunized with the indicated construct on Day 0 and boosted at week 4 (W4). Blood samples were taken at week 2 (W2) and week 6 (W6). **(B)** Antibody binding titers measured by ELISA. Plates were coated with the indicated construct and sera was added in serial ten-fold dilutions. Results are shown as EC50, calculated as –Log10 of the dilution that achieves 50% maximal binding. Each point represents sera from an individual mouse and means of each group are represented by an orange line. Statistical significance was determined by one-way ANOVA test followed by Tukey’s multiple comparison, and is represented by ** = p<0.01. **(C)** Neutralization of rVSV-SARS2_G614_Δ19 pseudovirus, given as IC50 values demarcating dilution at which 50% maximal neutralization is achieved. The LOD line represents the limit of detection of the experiment. Means and statistical significance are presented as shown in (B). **(D and E)** Neutralization of SARS-CoV-2 authentic viruses. PRNT50 titers with the authentic Wuhan_D614G SARS-CoV-2 **(D)** and the B1.351 lineage of SARS-CoV-2 (commonly referred to as the South Africa variant) **(E)**.

## DISCUSSION

The SARS-CoV2 spike (S) protein, in its pre-fusion assembly on the virion surface, is the main target of neutralizing and protective antibodies. A stabilized spike that maintains a pre-fusion conformation and trimeric assembly, as well as the natural RBD dynamics, is essential for development of effective vaccine candidates and research reagents. Such a molecule would ensure that the native quaternary structure is maintained to allow binding and elicitation of conformation-dependent neutralizing antibodies. Most vaccine candidates now in use or in clinical trials employ derivatives of the prototypical “2P” spike that carries two prolines. However, thermostability and yield of S-2P are both low. Improvements in yield, thermostability and quaternary structure of expressed spike protein would likely improve immunogenicity, efficacy, manufacturability and distribution of vaccines.

A second-generation spike construct, HexaPro, carries four additional prolines and offers greater yield and stability relative to S-2P. One proline substitution in both S-2P and HexaPro is K986P. The wild-type K986, however, forms salt bridge interactions that afford stability to the pre-fusion structure; for example, K986-D748 stabilizes the S2 core and K486-D427 is important for initial RBD positioning in the down position (Juraszek et al. 2021; Cai et al. 2020). In this study, we also show that HexaPro possesses some glycosylation patterns on S2 that are inconsistent with the authentic virus. Other versions of engineered spike introduce a disulfide bond to anchor the RBDs in the down position. This anchoring does not permit elicitation of, or binding by those antibodies that target the inner and upper surfaces of the RBD that are exposed upon transition to the up conformation. These various structural features and altered glycosylation together may alter immunogenicity and limit utility of early generation spikes.

In this work, we generated a spike antigen and vaccine candidate, VFLIP, which more effectively recapitulates the structural and antigenic properties of the native spike. Modifications in the VFLIP spike include: (i) a short flexible linker that replaces the S1/S2 cleavage fusion loop to prevent loss of S1 and springing of S2 to its post-fusion conformation; (ii) restoration of native Lys at position 986 to preserve the stabilizing K986-mediated salt bridges; and (iii) a disulfide bond at the base of spike between the S2 of one protomer and the S2’ of the adjacent protomer to maintain native trimerization without exogenous motifs.

The resulting VFLIP spike expresses to levels up to ten-fold higher than S-2P in both HEK293F and ExpiCHO cells, yielding abundant highly pure protein. This robust expression can facilitate production of recombinant protein vaccines. Further, more stable presentation of spike having a structure similar to that displayed on the virion will improve activity and possibly reduce the dose needed. Increased expression and a more stable encoded spike may also improve mRNA or viral-vectored vaccines by allowing higher expression and better antigenic presentation per nucleic acid molecule (Hsieh et al. 2020). These improvements could lower the dose of nucleic acid necessary to achieve robust in vivo expression and protection, reduce cost, lower exposure to reactogenic vaccine components, and broaden vaccine distribution.

Removing exogenous stabilizing domains prior to vaccine administration is also useful for preventing deleterious, and potentially confounding, immune responses. In December 2020, phase I/II clinical trials for the Australian COVID-19 vaccine, UQ-CSL V451, were halted due to false positives on some HIV-1 serology tests that were likely associated with the inclusion of a C-terminal “molecular clamp” derived from the HIV-1 env in the SARS-COV-2 spike (Update on UQ COVID-19 vaccine). An alternate trimerization domain, Foldon, is also highly immunogenic and may influence the resulting immune response (Brouwer et al. 2020; Lainšček et al. 2020).

With a 3 °C higher Tm, VFLIP is more thermostable than HexaPro and retains its trimeric structure even after removal of the Foldon trimerization domain (VFLIPΔFoldon). VFLIPΔFoldon remains trimeric after lyophilization, multiple freeze/thaw cycles, and prolonged storage at either 4 °C or at room temperature. Storage of VFLIPΔFoldon antigen in a drawer at room temperature for over a month has no detectable effect on spike quality. In contrast, HexaProΔFoldon is more prone to dissociate into monomers, with dissociation exacerbated by lyophilization, multiple freeze/thaw cycles, or prolonged storage at room temperature or 4 °C (Figure 5A-C). VFLIPΔFoldon also exhibits good immunogenicity in mice. Together these results suggest that trimerization domains can be removed without substantial impact on antigenic function.

Another important goal for antigen engineering is maintenance of native post-translational modifications, especially glycosylation, which plays a significant role in protein folding and stability. Glycans also contribute to the antigenic landscape by occluding regions of the protein from antibody access in addition to comprising portions of antibody epitopes (Grant et al. 2020; Walls, Park, et al. 2020). Therefore, accurate reflection of the glycan profile on authentic virions is critical for ensuring faithful mimicry of native antigenicity and immunogenicity. The VFLIP spike incorporates and displays native-like glycosylation that accurately reflects the glycan profile of the native virion (Yao et al. 2020). In both VFLIP and authentic virus, complex glycans are present at N603 and N709 in the conserved S2 subunit, whereas 5P and S-2P spike instead display high-mannose structures. VFLIP also has native-like processing of RBD glycans (N331 and N343) and NTD glycans, which have been shown to influence the conformational state of RBD (N165 and N234).

For practical utility as a vaccine or research reagent, a spike immunogen must be able to be efficiently produced, properly folded, and natively glycosylated in cell lines appropriate for industrial production (Blundell et al. 2020; Raska et al. 2010). ExpiCHO-S is a high-yield mammalian culture system derived from the commonly used industrial cell line, CHO-S, approved for Good-Manufacturing-Practice (GMP) production (He et al. 2018). The high yields of VFLIP spike obtained from the ExpiCHO expression system in our academic laboratory suggest that VFLIP would be amenable for large-scale industrial production. The VFLIP spike also better reflects the glycosylation of spikes present on the native virion, whether expressed in HEK293F or ExpiCHO cells (Yao et al. 2020).

The third-generation design of VFLIP preserves the natural RBD motion present in the native virion. High-resolution cryo-EM demonstrates that VFLIP displays the RBDs in a predominately “closed” conformation until interaction with the ACE2 receptor lifts the RBDs to the “up” conformation. In contrast, other engineered spikes employing disulfide bonds have all RBDs “locked down” and often positioned deeper than on the native virion. These structural perturbations may potentially affect the antigenicity of these designs: immunogens that begin with RBD “down” may better elicit those antibodies that benefit from an avidity boost and a symmetrical presentation of epitopes for B cell receptor cross-linking. Consistent with this possibility, an iso-affinity plot shows that CoVIC antibodies bound to a disulfide-bound “locked-down” spike (DS2) consistently display slower on-rates and lower overall affinity compared to VFLIP or HexaPro (Figure 4A).

For structural biology, VFLIP also enables extensive characterization of ACE2 and those antibodies that normally disrupt the spike structure through receptor mimicry and post-fusion triggering. Although methods such as brief incubation and immediate freezing have been attempted, structures of these complexes have remained elusive due to low site occupancy and fewer quality particles for high-resolution structural determination. In contrast, VFLIP exhibits slow off-rates for bound ACE2 or antibodies and is not disrupted by receptor-mimicking antibodies, thus increasing the number of high-quality particles with fully occupied epitopes and facilitating acquisition of structures.

Finally, we sought to determine if the innovative VFLIP design can serve as an effective immunogen for vaccination against SARS-CoV-2. The need for improved vaccine candidates is underscored by the recent emergence of variants of concern (VOC) that exhibit increased transmissibility and reduced sensitivity to neutralization by polyclonal sera and some therapeutic mAbs (D. Zhou et al. 2021; Planas et al. 2021; Wang et al. 2021). Lineage B.1.351 (known informally as the South African variant), has the greatest potential for antibody escape, largely mediated by the N501Y, E484K, and K417N substitutions in the RBD. Studies using pseudotyped and authentic B.1.351 have demonstrated a 6-14 fold reduction in neutralization potency of sera from COVID-19 patients or vaccinated individuals, and complete ablation of neutralization in up to 40% of convalescent samples (Planas et al. 2021; D. Zhou et al. 2021; Wang et al. 2021).

The high potency and cross-reactivity of VFLIP-induced sera is likely the result of the predominant “all-down” initial conformation and increased structural stability. Specifically, the more closed conformation of the VFLIP spike may dampen antibody responses against immunodominant epitopes on “up” RBDs that enable viral escape, including the variable RBM containing N501Y, E484K and K417N (Duan et al. 2020). Furthermore, the improved stability of the VFLIP spike trimer could improve generation of neutralizing antibodies. The structural stability of a vaccine antigen is reported to correlate with the development of highly matured antibodies that require temporally extended germinal center reactions (Feng et al. 2016; Karlsson Hedestam et al. 2017) (Cirelli and Crotty 2017). Stabilizing these spike proteins permits the extended presentation of native antigen on follicular dendritic cells (FDCs)–including conformational epitopes that are lost upon antigen degradation–while avoiding presentation of potentially distracting, non-native or undesired epitopes such as those present on inner surfaces that are only exposed in the monomer (Finney et al. 2018). Moreover, the resistance of VFLIP to fusogenic triggering prevents destruction of the antigen upon ligation of receptor-mimicking antibodies, which could extend the half-life of the antigen and, in turn, enhance generation of potently neutralizing antibodies (El Shikh et al. 2010; Doria-Rose and Joyce 2015; McLellan et al. 2013). In immunized BALB/c mice, VFLIP spikes elicited neutralizing titers superior to S-2P, and offers greater stability, and quaternary epitope presentation than HexaPro. This improved stability, and incorporation of more native glycan structures improves preservation and display of epitopes throughout antigen trafficking in vivo. Taken together, VFLIP is a highly stable, covalently-linked SARS-CoV-2 spike with improved structural and immunological properties, which can serve as an improved research tool and promising vaccine candidate for further study.

## ACKNOWLEDGEMENTS

We acknowledge support from National Institute of Health grant NIAID U19 AI 109762-S2 and the COVID-19 Therapeutics Accelerator of the Bill and Melinda Gates Foundation, Mastercard, Wellcome Trust and private philanthropic support to the CoVIC Consortium (EOS, SS, DB), Dutch Research Council NWO Gravitation 2013 BOO (JS), Institute for Chemical Immunology (ICI; 024.002.009; JS), the Medical Research Future Fund 2020 Antiviral Development Call (2001739 to DC), Bill and Melinda Gates Foundation INV-004923 (I.A.W.), Fellowship 1157744 from the National Health and Medical Research Council and Fellowship DE190100985 from the Australian Research Council (RR), and National Key Research and Development Project (2020YFC0848500)

## AUTHOR CONTRIBUTIONS

Conceptualization, E.O, and E.O.S.; Methodology, E.O., C.M., J.S., W.P., Y.W. and E.O.S.; Investigation E.O., C.M., W.P., Y.W., M.Y.; Formal Analysis, E.O., C.M., J.S., W.P., Y.W. and E.O.S.; Writing – Original Draft, E.O., C.M., E.O.S.; Writing – Review & Editing, E.O., C.M., E.O.S., S.S., J.S., Visualization, Supervision, E.O.S., J.S..; Resources, G.l., R.R., D.C., W.J; Funding Acquisition, E.O.S., J.S., D.C., W.J., T.G., I.A.W.

## DECLARATION OF INTERESTS

The authors declare no competing interests.

## SUPPLEMENTAL ITEM LEGENDS

**Extended Data Table S1. Summary of SARS-CoV-2 spike variants used in this study. Related to** Figure 1.

List of spike constructs and their features including proline backbone used (S-2P PDB: 6VSB, 6P PBD: 6XKL, 5P new in this study), S1/S2 linker sequence (if present), cysteine mutations for formation of disulfide bonds and the relative yield using the area under the curve of the size-exclusion trimer peak for the constructs expressed in HEK293F and ExpiCHO cell lines.

**Extended Data Table S2. High-throughput surface plasmon resonance measurement of antibody association kinetics. Related to** Figure 4.

The association and dissociation rates for each antibody (referenced using the COVIC ID) were used to calculate the affinity for soluble RBD (green), HexaPro (blue), VFLIP (red) and DS2 (yellow). Cells colored yellow are flagged as a poor fit, with the standard deviation of the residuals being > 10% of the calculated R_max_ value. Purple shading indicates that the fitting logic reached one of the set limits. N/A= not assessed.

**Extended Data Fig S1. CryoEM characterization of VFLIP. Related to** Figure 2.

**(A)** The op portion of the panel shows a representative micrograph of VFLIP spike. The bottom portion shows ten reference-free 2D class averages. **(B)** FSC curve and viewing distribution plot generated in cryoSPARC V2.15 for VFLIP reconstruction. **(C)** Cryo-EM density colored according to local resolution.

**Extended Data Fig S2. CryoEM characterization of VFLIP_G614 variant. Related to** Figure 2.

**(A)** Top portion of the panel shows a representative micrograph of VFLIP spike G614 variant. Bottom portion shows ten reference-free 2D class averages. **(B)** Side and top views of the electron density map for VFLIP D614G variant. Each protomer surface is shown in either pink, green or blue, and N-linked glycan densities are shown orange. **(C)** FSC curve and viewing distribution plot generated in cryoSPARC V2.15 for VFLIP_D614G reconstruction. **(D)** Cryo-EM density is colored according to local resolution.

**Extended Data Fig S3. Restoration of the K986-D748 salt-bridge in 5P spikes. Related to** Figure 2. Side view of the electron density map for the VFLIP Wuhan variant is shown in transparent gray. One protomer in ribbon is shown in cyan. The expanded view shows a section of the monomer with the density indicating formation of a salt bridge between K986-D748. Atomic interactions within the amino acids are shown with dashed yellow lines and estimated distances.

**Extended Data Fig S4. LC-MS/MS analysis of S oligosaccharides from 5P and VFLIP expressed in HEK293F cells and VFLIP expressed in ExpiCHO cells. Related to** Figure 3. The fraction of glycan species present on 5P and VFLIP produced in HEK293F cells (orange and cyan, respectively) and VFLIP produced in ExpiCHO cells (purple) is shown for the indicated asparagine residue. Green, yellow and orange lines at the top of each plot represent high-mannose, hybrid and complex glycans, respectively.

**Extended Data Fig S5. CryoEM characterization of VFLIP_G614 variant in complex with HLX70 Fab fragment. Related to** Figure 4.

**(A)** A representative micrograph of VFLIP_G614 in complex with HLX70 is shown in the top portion of the panel, while the bottom portion shows ten reference-free 2D class averages. **(B)** Fitting of the cryo-EM density map of VFLIP_D614G in complex with the Fab portion of HLX70 using PDB ID 7CWM. **(C)** FSC curve generated in cryoSPARC V2.15 for the reconstruction of VFLIP_D614G in a complex with three HLX70 ACE2-Fc. **(D)** Biolayer Interferometry analysis of HLX70 mAb binding kinetics with VFLIP immobilized on the chip. Multiple concentrations of the ACE2-Fc were evaluated and are shown on the legend. Global fit curves are shown as black dotted lines.

**Extended Data Fig S6. CryoEM characterization of VFLIP_G614 glycoprotein in complex with the HLX71 ACE2-Fc. Related to** Figure 4.

**(A)** A representative micrograph of VFLIP_G614 in complex with HLX71 is shown in the top portion of the panel, while the bottom portion shows ten reference-free 2D class averages. **(B)** Fitting of the cryo-EM density map of VFLIP_D614G in complex with the Fab portion of HLX71 usingPDB ID: 7A98. **(C)** FSC curve generated in cryoSPARC V2.15 for the reconstruction of VFLIP_D614G in a complex with three HLX71 Fab fragments. **(D)** Biolayer Interferometry analysis of HLX71 ACE2-Fc binding kinetics with VFLIP immobilized on the chip. Multiple concentrations of the HLX71 Ab were monitored and are shown on the legend. Global fit curves are shown as black dotted lines.

**Extended Data Fig S6. Related to** Figure 5. Peak fraction analysis of HexaProΔFoldon and VFLIPΔFoldon complexes shown in Figure 5 of the main text using SDS-PAGE with (D) and without (E) glutaraldehyde cross-linking.

**Extended Data Fig S7. Extension of VFLIP modifications to SARS-CoV-1 spike. Related to Discussion.** Adaptation of VFLIP technology to another β-Coronavirus. (A) Expression of SARS1_S-2P and SARS1_VFLIP constructs produced from 50 ml ExpiCHO cells. (B) Non-reducing SDS-PAGE in the presence (+) or absence (-) of glutaraldehyde cross-linking. In the non-cross-linked condition, VFLIP showed a well-formed inter-monomeric disulfide bond in positions homologous to those on SARS-CoV-2 VFLIP spike.

## STAR METHODS

### RESOURCE AVAILABILITY

#### Lead Contact

Further information and requests for resources and reagents should be directed to and will be fulfilled by the Lead Contact, Erica Ollmann Saphire.

#### Materials Availability

All reagents generated in this study are available from the Lead Contact upon request, but we may require a completed material transfer agreement

#### Data and Code Availability

Datasets generated during this study are included in the article or are available from the corresponding authors on request. The cryo-EM maps generated during this study are deposited in the Electron Microscopy Data Bank EMBL-EBI under accession codes EMD-23886, EMD-23889, EMD-23891, and EMD-23896.

The atomic coordinates generated during this study are deposited in the RCSB Protein Data Bank.

The LC-MS/MS glycoproteomics data have been deposited in the ProteomeXchange Consortium via the PRIDE partner repository under the dataset identifier PXD025554.

### EXPERIMENTAL MODEL AND SUBJECT DETAILS

#### Bacterial strains

*E. coli* strain Rosetta DE3 (Novagen) was grown in lysogeny broth. The genotype is: F-ompT hsdSB(rB-mB-) gal dcm (DE3) pRARE (CamR). Selection markers were used at the indicated concentrations: ampicillin (100 μg/mL); chloramphenicol (28.3 μg/mL).

#### Cell lines

HEK-293T, Vero E6 and Vero-CCL81 cell lines were obtained from ATCC and cultured in DMEM medium (Gibco 31966021) supplemented with 10% Fetal Bovine Serum and incubated at 37 °C and 5% CO2. HEK-293F and Expi-CHO cells were obtained from Thermo Fisher Scientific and maintained in Expi293 Expression Medium and ExpiCHO-Expression Medium (Thermo Fisher Scientific), respectively.

## EXPERIMENTAL MODEL AND SUBJECT DETAILS

### METHOD DETAILS

#### Design of SARS-CoV-2 spike variants

Spike variants were initially designed using the S-2P construct that includes ectodomain residues 13-1208 (Genbank: MN908947), two proline substitutions (K986P, V967P), and substitution of cleavage site residues RRAR with GSAS at position 682-685 (“RRAR” to “GSAS”). Designs containing five or six prolines were based on the HexaPro construct that carries, in addition to the proline substitutions of S-2P, proline substitutions at positions 817, 892, 899 and 942. All variants were cloned into a PhCMV mammalian expression vector containing a C-terminal foldon trimerization domain followed by an HRV-3C cleavage site and a Twin-Strep-Tag. Using the published S-2P cryo-EM structure (PDB: 6VSB), truncated cleavage site linkers of varying length and flexibility were designed to prevent S1/S2 cleavage and improve protein expression. Candidate interprotomer disulfide bond candidates were selected by assessing residues with CB atoms lying within 5 Å of subunit interfaces, or by visual inspection. HexaPro P986 was reverted to lysine to restore a potential interprotomer salt-bridge that is disrupted by this mutation (PDB: 6VXX). Combinatorial variants containing different cleavage site linkers, proline substitutions and disulfide bonds were evaluated for effects on purity, yield and thermostability.

### Transient transfection and protein purification

SARS-CoV-2 spike variants were transiently transfected in Freestyle 293-F and ExpiCHO-S cells (Thermo Fisher). Both cell lines were maintained and transfected according to the manufacturer’s protocols. Briefly, 293-F cells were transfected with plasmid DNA mixed with polyethylenimine and harvested on day 5. Cultures were clarified by centrifugation, followed by addition of BioLock (IBA Life Sciences), passage through a 0.22 µM sterile filter, and purification on an ӒKTA go system (Cytivia) using a 5mL StrepTrap-HP column equilibrated with TBS buffer (25mM Tris pH 7.6, 200mM NaCl, 0.02% NaN_3_), and eluted in TBS buffer supplemented with 5mM d-desthiobiotin (Sigma Aldrich). Proteins were then purified by size-exclusion-chromatography (SEC) on a Superdex 6 Increase 10/300 column (Cytivia) in the same TBS buffer.

For all ExpiCHO cultures, the manufacturer’s “High Titer” protocol was used with a 7-day culture incubation to assess relative expression. Briefly, plasmid DNA and Expifectamine were mixed in Opti-PRO SFM (Gibco) according to the manufacturer’s instructions, and added to the cells. On day 1, cells were fed with manufacturer-supplied feed and enhancer as specified in the manufacturer’s protocol, and cultures were moved to a shaker incubator set to 32 °C, 5% CO_2_ and 115 RPM. On day 7, the cultures were clarified by centrifugation, BioLock was added, and supernatants were passed through a 0.22 µM sterile filter. Purification was performed as above, on an ӒKTA go system using a 5mL StrepTrap HP column and TBS buffer, followed by SEC on a Superdex 6 Increase 10/300 column with TBS buffer.

### Differential Scanning Calorimetry

Thermal stability was measured using a MicroCal VP-Capillary calorimeter (Malvern) with 0.6 mg/ml of each sample in phosphate-buffered saline (PBS) buffer at a scanning rate of 90 °C hour−1 from 20°C to 120°C. Data were analyzed using VP-Capillary DSC automated data analysis software.

### Negative stain electron microscopy (NS-EM)

To prepare samples for NS-EM, 3 µL 0.02 mg/mL protein was applied to a carbon film 400 mesh grid for 1 minute. The grid was then washed three times with 10 µL Milli-Q water and stained three times with 4 µL drops of 1% uranyl formate, with the first two drops briefly and the third time for 1 minute. The grid was blotted with Whatman filter paper after each application of liquid. Micrographs were collected on a Titan Halo transmission electron microscope, operating with an accelerating voltage of 300kV and using a pixel size of 1.4Å/pixel.

### Cryo-Electron Microscopy sample preparation

VFLIP and VFLIP_D614G were concentrated to 2mg/ml and electron microscopy grids were prepared by placing a 3 µL aliquot of the sample on a plasma-cleaned C-flat grid (2/1C-3T, Protochips Inc) that was then immersed in liquid ethane for vitrification. For formation of VFLIP_D614G:HLX70 Fab and VFLIP_D614G:HLX71_ACE2-Fc complexes, the trimerization tag ‘Foldon’ and purification tags of VFLIP_D614G spike were enzymatically removed by an overnight treatment with HRV protease at room temperature before concentration to 1mg/ml and incubation at a 1:2 molar ratio with HLX70 Fab or HLX71 ACE2-Fc at room temperature overnight. The samples were then injected over a gel filtration column (Superose 6 10/30, GE Life Sciences) equilibrated with 20mM Tris pH 8.0 and 150mM NaCl. The complex peak fractions were concentrated to an absorbance of 2.0. Electron microscopy grids were prepared as described above.

### Cryo-EM data collection and processing

Grids were loaded into a Titan Krios G3 electron microscope (Thermo Fisher Scientific) equipped with a K3 direct electron detector (Gatan, Inc.) at the end of a BioQuantum energy filter, using an energy slit of 20eV. The microscope was operated with an accelerating voltage of 300kV. Grids were imaged with a pixel size of 0.66 Å in counting mode. Data was acquired using the software EPU. Motion correction, CTF estimation, and particle-picking were done with Warp (Tegunov and Cramer 2019). Extracted particles were exported to cryoSPARC-v2 (Punjani et al. 2017) (Structura Biotechnology Inc.) for 2D classification, ab initio 3D reconstruction, and refinement. C1 symmetry was used during homogeneous refinement.

### Cryo-EM model building and analysis

A previously published structure of the SARS-CoV-2 ectodomain with all RBDs in the down conformation (PDB ID 6X79) was used to fit the cryo-EM maps in UCSF ChimeraX (Goddard et al. 2018) and PyMOL. Mutations were made in PyMOL. Coordinates were then fitted manually using COOT (Emsley et al. 2010), followed by cycles of refinement using Phenix (Afonine et al. 2018) real space refinement. COOT was used for subsequent fitting.

### Glycoproteomics sample preparation

Recombinant SARS-CoV-2 spike protein was denatured at 95 °C at a final concentration of 2% sodium deoxycholate (SDC), 200 mM Tris/HCl, 10 mM tris(2-carboxyethyl)phosphine, pH 8.0 for 10 min, followed by a 30 min reduction at 37 °C. Next, samples were alkylated by adding 40 mM iodoacetamide and incubated in the dark at room temperature for 45 min. For each protease digestion, 3 μg recombinant SARS-CoV-2 spike protein was used. Samples were divided in thirds for parallel digestion with gluC (Sigma)-trypsin (Promega), chymotrypsin (Sigma) and alpha lytic protease (Sigma). For each protease digestion, 18 μL of the denatured, reduced, and alkylated samples was diluted in a total volume of 100 μL 50 mM ammonium bicarbonate and proteases were added at a 1:30 ratio (w:w) for incubation overnight at 37 °C. For the gluC-trypsin digestion, gluC was added first for two hours, and then incubated with trypsin overnight. After overnight digestion, SDC was removed by precipitation with2 μL formic acid and centrifugation at 14,000 rpm for 20 min. The resulting supernatant containing the peptides was collected for desalting on a 30 µm Oasis HLB 96-well plate (Waters). The Oasis HLB sorbent was activated with 100% acetonitrile and subsequently equilibrated with 10% formic acid in water. Next, peptides were bound to the sorbent, washed twice with 10% formic acid in water and eluted with 100 µL 50% acetonitrile/10% formic acid in water (v/v). The eluted peptides were dried under vacuum and resuspended in 100 µL 2% formic acid in water. The experiment was performed in duplicate.

### Glycoproteomics mass spectrometry

The duplicate samples were analyzed with two different mass spectrometry methods, using identical LC-MS parameters andt distinct fragmentation schemes. In one method, peptides were subjected to Electron Transfer/Higher-Energy Collision Dissociation fragmentation (Frese et al. 2012, 2013). In the other method, all precursors were subjected to HCD fragmentation, with additional EThcD fragmentation triggered by the presence of glycan reporter oxonium ions. For each duplicate sample injection, approximately 0.15 μg of peptides were run on an Orbitrap Fusion Tribrid mass spectrometer (Thermo Fisher Scientific, Bremen) coupled to a Dionex UltiMate 3000 (Thermo Fisher Scientific). A 90-min LC gradient from 0% to 44% acetonitrile was used to separate peptides at a flow rate of 300 nl/min. Peptides were separated using a Poroshell 120 EC-C18 2.7-Micron analytical column (ZORBAX Chromatographic Packing, Agilent) and a C18 PepMap 100 trap column (5 mm x 300 µm, 5 µm, Thermo Fisher Scientific). Data was acquired in data-dependent mode. Orbitrap Fusion parameters for the full scan MS spectra were as follows: a standard AGC target at 60 000 resolution, scan range 350-2000 m/z, Orbitrap maximum injection time 50 ms. The ten most intense ions (2+ to 8+ ions) were subjected to fragmentation. For the EThcD fragmentation scheme, the supplemental higher energy collision dissociation energy was set at 27%. MS2 spectra were acquired at a resolution of 30,000 with an AGC target of 800%, maximum injection time 250 ms, scan range 120-4000 m/z and dynamic exclusion of 16 s. For the triggered HCD-EThcD method, the LC gradient and MS1 scan parameters were identical. The ten most intense ions (2+ to 8+) were subjected to HCD fragmentation with 30% normalized collision energy from 120-4000 m/z at 30,000 resolution with an AGC target of 100% and a dynamic exclusion window of 16 s. Scans containing any of the following oxonium ions within 20 ppm were followed up with additional EThcD fragmentation with 27% supplemental HCD fragmentation. The triggering reporter ions were: Hex(1) (129.039; 145.0495; 163.0601), PHex(1) (243.0264; 405.0793), HexNAc(1) (138.055; 168.0655; 186.0761), Neu5Ac(1) (274.0921; 292.1027), Hex(1)HexNAc(1) (366.1395), HexNAc(2) (407.166), dHex(1)Hex(1)HexNAc(1) (512.1974), and Hex(1)HexNAc(1)Neu5Ac(1) (657.2349). EThcD spectra were acquired at a resolution of 30,000 with a normalized AGC target of 400%, maximum injection time 250 ms, and scan range 120-4000 m/z.

### Mass spectrometry data analysis

The acquired data was analyzed using Byonic (v3.11.1) against a custom database of SARS-CoV-2 spike protein sequences and the proteases used in the experiment to search for glycan modifications with 12/24 ppm search windows for MS1 and MS2, respectively. Up to five missed cleavages were permitted using C-terminal cleavage at R/K/E/D for gluC-trypsin or F/Y/W/M/L for chymotrypsin. Up to 8 missed cleavages were permitted using C-terminal cleavage at T/A/S/V for alpha lytic protease. Carbamidomethylation of cysteine was set as a fixed modification and oxidation of methionine/tryptophan was set as variable rare 1. N-glycan modifications were set as variable common 2, allowing up to a maximum of 3 variable common and 1 rare modification per peptide. All N-linked glycan databases from Byonic were merged into a single non-redundant list for inclusion in the database search. All reported glycopeptides in the Byonic result files were first filtered for score ≥ 100 and PEP2D ≤ 0.01, then manually inspected for quality of fragment assignments. All glycopeptide identifications were merged into a single non-redundant list per sequon. Glycans were classified based on HexNAc and Hexose content as paucimannose (2 HexNAc, 3 Hex), high-mannose (2 HexNAc; > 3 Hex), hybrid (3 HexNAc) or complex (> 3 HexNAc). Byonic search results were exported into mzIdentML format to build a spectral library in Skyline (v20.1.0.31) and to extract peak areas for individual glycoforms from MS1 scans. N-linked glycan modifications identified from Byonic were manually added to the Skyline project file in XML format. Reported peak areas were pooled based on the number of HexNAc, Fuc or NeuAc residues to distinguish paucimannose, high-mannose, hybrid, and complex glycosylation.

### CoVIC antibodies

Eukaryotic expression vectors containing the DNA sequence of HLX70 and HLX71 were used to stably transfect CHO cells. Clinical-grade recombinant human antibodies produced from CHO cells by Shanghai Henlius Biotech, Inc. according to Good Manufacturing Practice guidelines and formulated as a preservative-free lyophilized powder.

### High-throughput surface plasmon resonance

High-throughput SPR capture kinetic experiments were performed on an LSA biosensor system equipped with a planar carboxymethyldextran CMDP sensor chip (Carterra). The LSA automates the choreography between two microfluidic modules, namely a single flow cell (SFC), which flows samples over the entire array surface and a 96-channel printhead (96PH) used to create arrays of up 384 samples. The capture surface was prepared using the SFC by standard amine-coupling of goat anti-human IgG Fc (Southern Biotech) to create a uniform surface, or lawn, over the entire chip. The system running buffer was 1X HBSTE (10 mM HEPES pH 7.4, 150 mM NaCl, 3mM EDTA, 0.05% Tween-20). The chip was activated with a 10-minute injection of freshly prepared 1:1:1 (v/v/v) 0.4M EDC 0.1MN-hydroxysulfosuccinimide (SNHS) with 0.1M 2-(N-morpholino)ethanesulfonic acid (MES) pH 5.5 before coupling of goat anti-human IgG Fc (50 μg/ml in 10 mM sodium acetate pH 4.5) for 15 minutes. Excess reactive esters were blocked with a 7-minute injection of 1M ethanolamine HCl pH 8.5. Final coupled levels (mean ± Std.Dev. RU across all 384 array regions of interest (ROIs) were 535 ± 32RU. After preparing the capture surface, the instrument was primed using assay running buffer (HBSTE with 0.5 mg/mL BSA). The Fc-ligands and mAbs were diluted into assay running buffer and captured onto the array using the 96PH for 15 minutes at three dilutions of 25, 3.6, and 0.9. Antibodies were captured and buffer blanks were then injected followed by a titration series of increasing antigen concentration. RBD and spike proteins were injected at 0.8, 2.5, 7.4, 22, 67, and 200 nM for 5 minutes with a 15-minute dissociation. After each antigen titration series the surface was regenerated with three, 60-second pulses of 0.475% H_3_PO_4_. Binding data from the local reference spots (interspots, representing the unoccupied capture surface) were subtracted from the active ROIs and the nearest buffer blank analyte responses were subtracted to double-reference the data. The double-referenced data were fit globally to a simple 1:1 Langmuir binding model using the Carterra Kinetics software tool to provide *k*a, *k*d, and Rmax values for each spot.

### Biophysical experiments and spike-splitting tests of VFLIPΔFoldon

Enzymatic removal of the ‘Foldon’ trimerization tag from VFLIP and HexaPro was facilitated by cloning the HRV-C3 cleavage site followed by Strep purification tags between the C terminus of the SARS-CoV-2 spike and the Foldon. After purification on a StrepTrap HP column, proteins were incubated overnight at room temperature with 2U HRV-3C protease per 100 μg protein at room temperature. The cleaved proteins were then SEC purified. For biophysical characterization, 150 μg of each protein was incubated at 4 °C and 37 °C for 5 days and then SEC purified. The same amount of protein was subjected to 10 cycles of fast freeze/thaw and then SEC purified. For lyophilization, 150 μg Foldon-free spike was dehydrated overnight using a SpeedVac RT and the lyophilized proteins were resuspended 5 days later in TBS before SEC purification.

To form immune complexes between shACE2 and B6 Fab fragments, and offer the greatest chance for spike separation as we observed for these molecules with other forms of spike, we incubated complexes for 2 days at 4°C at a 1:2 molar ratio with Foldon-free proteins concentrated to 1 mg/ml. Samples were purified by SEC as described above.

### Mice immunization

For mouse immunization and serum extraction Institutional Animal Care and Use Committee (IACUC) guidelines were followed with animal subjects tested in the immunogenicity study. Six-week-old BALB/c mice were purchased from the Jackson Laboratory. The mice were housed in ventilated cages in environmentally controlled rooms at the LJI animal facility, in compliance with an approved IACUC protocol and AAALAC (Association for Assessment and Accreditation of Laboratory Animal Care) International guidelines. At week 0, each mouse was immunized with 25 µg of the indicated antigen in 50 µl PBS together 25 μl of the Magic Mouse CpG adjuvant (Creative technologies) and 25 μl aluminum hydroxide (Invivogen) administered by an intramuscular (i.m.) route. At week 4, the animals were boosted with the same antigen/adjuvant composition as used for the prime. At week 6, the animals were bled through the retro-orbital membrane using fractionator tubes. Sera were heat inactivated at 56 °C for 1 hour and stored at −80 °C until analysis.

### Enzyme-linked immunosorbent assays

96-well EIA/RIA plates (Corning, Sigma) were coated with 0.1 μg per well of HexaPro in PBS and incubated at 4 °C overnight. On the following day, the coating solution was removed and wells were blocked with 5% skim milk diluted in PBS with 0.1% Tween 20 (PBST) at room temperature for 1 h. Mouse serum samples that had been previously heat inactivated at 56 °C for 1 h were diluted 1:50 and the serially diluted five-fold in 5% skim milk in PBST. The blocking solution was removed and 50 μl of the diluted sera was added to the plates and incubated for 1 h at room temperature. Following incubation, the diluted sera were removed and the plates were washed 4 times with PBST. Goat anti-human IgG secondary antibody-peroxidase (Fc-specific, Sigma) diluted 1:3,000 in 5% skim milk in PBST was then added and the plates were incubated for 1 h at room temperature before washing four times with PBST. The ELISA was developed using 3,5,3′,5′-tetramethylbenzidine (Thermo Fisher Scientific) solution and the reaction was stopped after 5 min incubation with 4N sulfuric acid. The OD450 was measured using a Tecan Spark 10M plate reader. The dilution of each serum sample required to obtain a 50% maximum signal (EC50) against HexaPro was determined using nonlinear regression analysis in Prism 8 version 8.4.2 (GraphPad).

### rVSV SARS2 pseudovirus neutralization assay

Recombinant SARS-CoV-2-pseudotyped VSV-ΔG-GFP was generated by transfecting 293T cells with phCMV3-SARS-CoV-2 full-length spike carrying the D614G mutation and deletion of the 19 C-terminal amino acid using TransIT according to the manufacturer’s instructions. At 24 hr post-transfection, cells were washed 2x with OptiMEM and then infected with rVSV-G pseudotyped ΔG-GFP parent virus (VSV-G*ΔG-GFP) at MOI = 2 for 2 hours with rocking. The virus was then removed, and the cells were washed twice with OPTI-MEM containing 2% FBS (OPTI-2) before addition of fresh OPTI-2. Supernatants containing rVSV-SARS-2 pseudoviruses were removed 24 hours post-infection and clarified by centrifugation, pooled and stored at -−80 °C until use.

SARS-CoV-2-pseudotyped VSV-ΔG-GFP was next titered in Vero cells (ATCC CCL-81). Cells were seeded in 96-well plates at a sufficient density to form a monolayer at the time of infection. 10-fold serial dilutions of pseudovirus were made and added to cells in triplicate wells. Infection was allowed to proceed for 16-18 hr at 37 °C before fixation of the cells with 4% PFA and staining with Hoechst (10 µg/mL) in PBS. Fixative/stain was replaced with PBS and pseudovirus titers were quantified as the number of GFP-positive cells (fluorescent forming units, ffu/mL) using a CellInsight CX5 imager (Thermo Scientific) and automated enumeration of cells expressing GFP.

Mouse sera neutralization assays were performed with pre-titrated amounts of rVSV-SARS-CoV-2 pseudovirus with sera samples diluted 1:100 and serial four-fold dilutions. The pseudovirus and sera samples were incubated together at 37 °C for 1 hr before addition to confluent Vero monolayers in 96-well plates. The plates were incubated for 16-18 hrs at 37 °C in 5% CO_2_, and then the cells were fixed with4% paraformaldehyde and stained with 10 µg/mL Hoechst. Cells were imaged using a CellInsight CX5 imager and infection was quantified by automated enumeration of total cells and those expressing GFP. Infection was normalized to the average number of cells infected with rVSV-SARS-CoV-2 incubated without sera, and pooled sera from untreated mice was used as control. Data are presented as neutralization IC50 titers calculated using “One-Site Fit LogIC50” regression in GraphPad Prism 9.0.

### Plaque reduction neutralization (PRNT) assay

SARS-CoV-2 variant D614G was obtained through BEI Resources, NIAID, NIH: SARS-Related Coronavirus 2, Isolate Germany/BavPat1/2020, NR-52370. SARS-CoV-2 strain B1.351 was obtained through BEI Resources, NIAID, NIH: SARS-Related Coronavirus 2, Isolate hCoV-19/South Africa/KRISP-K005325/2020, NR-54009. Both SARS-CoV-2 D614G and B1.351 were propagated in Vero-CCL81 cells, titrated by plaque assay on Vero E6 cells, deep-sequenced by the La Jolla Institute for Immunology Sequencing Core. Assays were performed in the BSL3 facility at La Jolla Institute for Immunology. For PRNT assay, mouse serum was serial 5-fold diluted, starting from 50-fold to 156250-fold, before co-culture with 30-40 plaque forming units (PFU) of SARS-CoV-2 D614G or B1.351 for 1 h at 37 °C. The serum/virus mixture was then transferred onto Vero E6 cells (8 x 104 cells/well, 24-well plate) for 2 h at 37 °C. The inoculum was removed before overlaid with 1% carboxymethylcellulose medium to each well. All the conditions were tested in duplication. After 3 days cultivation, cells were fixed with 10 % formaldehyde in PBS for 30 min at RT prior stained with 0.1% crystal violet solution for 20 min at RT. Serum titer (NT50) was determined as the highest sample dilution that neutralize 50% of virus plaques.

## Notes

### Competing Interest Statement

The authors have declared no competing interest.

## REFERENCES

Afonine, Pavel V., Billy K. Poon, Randy J. Read, Oleg V. Sobolev, Thomas C. Terwilliger, Alexandre Urzhumtsev, and Paul D. Adams. 2018. “Real-Space Refinement in PHENIX for Cryo-EM and Crystallography.” Acta Crystallographica. Section D, Structural Biology 74 (Pt 6): 531–44.

Benton, Donald J., Antoni G. Wrobel, Pengqi Xu, Chloë Roustan, Stephen R. Martin, Peter B. Rosenthal, John J. Skehel, and Steven J. Gamblin. 2020. “Receptor Binding and Priming of the Spike Protein of SARS-CoV-2 for Membrane Fusion.” Nature 588 (7837): 327–30.

Blundell, Patricia A., Dongli Lu, Anne Dell, Stuart Haslam, and Richard J. Pleass. 2020. “Choice of Host Cell Line Is Essential for the Functional Glycosylation of the Fc Region of Human IgG1 Inhibitors of Influenza B Viruses.” Journal of Immunology 204 (4): 1022–34.

Bos, Rinke, Lucy Rutten, Joan E. M. van der Lubbe, Mark J. G. Bakkers, Gijs Hardenberg, Frank Wegmann, David Zuijdgeest, et al. 2020. “Ad26 Vector-Based COVID-19 Vaccine Encoding a Prefusion-Stabilized SARS-CoV-2 Spike Immunogen Induces Potent Humoral and Cellular Immune Responses.” NPJ Vaccines 5: 91.

Brouwer, Philip J. M., Tom G. Caniels, Karlijn van der Straten, Jonne L. Snitselaar, Yoann Aldon, Sandhya Bangaru, Jonathan L. Torres, et al. 2020. “Potent Neutralizing Antibodies from COVID-19 Patients Define Multiple Targets of Vulnerability.” Science 369 (6504): 643– 50.

Cai, Yongfei, Jun Zhang, Tianshu Xiao, Hanqin Peng, Sarah M. Sterling, Richard M. Walsh Jr, Shaun Rawson, Sophia Rits-Volloch, and Bing Chen. 2020. “Distinct Conformational States of SARS-CoV-2 Spike Protein.” Science 369 (6511): 1586–92.

Cao, Liwei, Jolene K. Diedrich, Daniel W. Kulp, Matthias Pauthner, Lin He, Sung-Kyu Robin Park, Devin Sok, et al. 2017. “Global Site-Specific N-Glycosylation Analysis of HIV Envelope Glycoprotein.” Nature Communications 8: 14954.

Casalino, Lorenzo, Zied Gaieb, Jory A. Goldsmith, Christy K. Hjorth, Abigail C. Dommer, Aoife M. Harbison, Carl A. Fogarty, et al. 2020. “Beyond Shielding: The Roles of Glycans in the SARS-CoV-2 Spike Protein.” ACS Central Science 6 (10): 1722–34.

Cirelli, Kimberly M., and Shane Crotty. 2017. “Germinal Center Enhancement by Extended Antigen Availability.” Current Opinion in Immunology 47: 64–69.

Corbett, Kizzmekia S., Darin K. Edwards, Sarah R. Leist, Olubukola M. Abiona, Seyhan Boyoglu-Barnum, Rebecca A. Gillespie, Sunny Himansu, et al. 2020. “SARS-CoV-2 mRNA Vaccine Design Enabled by Prototype Pathogen Preparedness.” Nature 586 (7830): 567– 71.

Doria-Rose, Nicole A., and M. Gordon Joyce. 2015. “Strategies to Guide the Antibody Affinity Maturation Process.” Current Opinion in Virology 11 (April): 137–47.

Duan, Liangwei, Qianqian Zheng, Hongxia Zhang, Yuna Niu, Yunwei Lou, and Hui Wang. 2020. “The SARS-CoV-2 Spike Glycoprotein Biosynthesis, Structure, Function, and Antigenicity: Implications for the Design of Spike-Based Vaccine Immunogens.” Frontiers in Immunology 11: 576622.

Edwards, Robert J., Katayoun Mansouri, Victoria Stalls, Kartik Manne, Brian Watts, Rob Parks, Sophie M. C. Gobeil, et al. 2021. “Cold Sensitivity of the SARS-CoV-2 Spike Ectodomain.” Nature Structural & Molecular Biology. 28, 128–131.

El Shikh, Mohey Eldin M., Rania M. El Sayed, Selvakumar Sukumar, Andras K. Szakal, and John G. Tew. 2010. “Activation of B Cells by Antigens on Follicular Dendritic Cells.” Trends in Immunology 31 (6): 205–11.

Emsley, P., B. Lohkamp, W. G. Scott, and K. Cowtan. 2010. “Features and Development of Coot.” Acta Crystallographica. Section D, Biological Crystallography 66 (Pt 4): 486–501.

“Evaluate Safety and Pharmacokinetics of HLX70 in Healthy Adult Volunteers.” n.d. Accessed April 30, 2021. https://clinicaltrials.gov/ct2/show/NCT04561076.

“Evaluate the Safety, Tolerability, Pharmacodynamics, Pharmacokinetics, and Immunogenicity of HLX71 (Recombinant Human Angiotensin-Converting Enzyme 2-Fc Fusion Protein for COVID-19) in Healthy Adult Subjects.” n.d. Accessed April 30, 2021. https://clinicaltrials.gov/ct2/show/NCT04583228.

Feng, Yu, Karen Tran, Shridhar Bale, Shailendra Kumar, Javier Guenaga, Richard Wilson, Natalia de Val, et al. 2016. “Thermostability of Well-Ordered HIV Spikes Correlates with the Elicitation of Autologous Tier 2 Neutralizing Antibodies.” PLoS Pathogens 12 (8): e1005767.

Finney, Joel, Chen-Hao Yeh, Garnett Kelsoe, and Masayuki Kuraoka. 2018. “Germinal Center Responses to Complex Antigens.” Immunological Reviews 284 (1): 42–50.

Frese, Christian K., A. F. Maarten Altelaar, Henk van den Toorn, Dirk Nolting, Jens Griep-Raming, Albert J. R. Heck, and Shabaz Mohammed. 2012. “Toward Full Peptide Sequence Coverage by Dual Fragmentation Combining Electron-Transfer and Higher-Energy Collision Dissociation Tandem Mass Spectrometry.” Analytical Chemistry 84 (22): 9668–73.

Frese, Christian K., Houjiang Zhou, Thomas Taus, A. F. Maarten Altelaar, Karl Mechtler, Albert J. R. Heck, and Shabaz Mohammed. 2013. “Unambiguous Phosphosite Localization Using Electron-Transfer/higher-Energy Collision Dissociation (EThcD).” Journal of Proteome Research 12 (3): 1520–25.

Ge, Jiwan, Ruoke Wang, Bin Ju, Qi Zhang, Jing Sun, Peng Chen, Senyan Zhang, et al. 2021. “Antibody Neutralization of SARS-CoV-2 through ACE2 Receptor Mimicry.” Nature Communications 12 (1): 250.

Gobeil, Sophie M-C, Katarzyna Janowska, Shana McDowell, Katayoun Mansouri, Robert Parks, Kartik Manne, Victoria Stalls, et al. 2021. “D614G Mutation Alters SARS-CoV-2 Spike Conformation and Enhances Protease Cleavage at the S1/S2 Junction.” Cell Reports 34 (2): 108630.

Goddard, Thomas D., Conrad C. Huang, Elaine C. Meng, Eric F. Pettersen, Gregory S. Couch, John H. Morris, and Thomas E. Ferrin. 2018. “UCSF ChimeraX: Meeting Modern Challenges in Visualization and Analysis.” Protein Science: A Publication of the Protein Society 27 (1): 14–25.

Grant, Oliver C., David Montgomery, Keigo Ito, and Robert J. Woods. 2020. “Analysis of the SARS-CoV-2 Spike Protein Glycan Shield Reveals Implications for Immune Recognition.” Scientific Reports 10 (1): 14991.

Güthe, Sarah, Larisa Kapinos, Andreas Möglich, Sebastian Meier, Stephan Grzesiek, and Thomas Kiefhaber. 2004. “Very Fast Folding and Association of a Trimerization Domain from Bacteriophage T4 Fibritin.” Journal of Molecular Biology 337 (4): 905–15.

Hastie, Kathryn M., Michelle A. Zandonatti, Lara M. Kleinfelter, Megan L. Heinrich, Megan M. Rowland, Kartik Chandran, Luis M. Branco, James E. Robinson, Robert F. Garry, and Erica Ollmann Saphire. 2017. “Structural Basis for Antibody-Mediated Neutralization of Lassa Virus.” Science 356 (6341): 923–28.

He, Linling, Sonu Kumar, Joel D. Allen, Deli Huang, Xiaohe Lin, Colin J. Mann, Karen L. Saye-Francisco, et al. 2018. “HIV-1 Vaccine Design through Minimizing Envelope Metastability.” Science Advances 4 (11): eaau6769.

Henderson, Rory, Robert J. Edwards, Katayoun Mansouri, Katarzyna Janowska, Victoria Stalls, Sophie M. C. Gobeil, Megan Kopp, et al. 2020. “Controlling the SARS-CoV-2 Spike Glycoprotein Conformation.” Nature Structural & Molecular Biology 27 (10): 925–33.

Hsieh, Ching-Lin, Jory A. Goldsmith, Jeffrey M. Schaub, Andrea M. DiVenere, Hung-Che Kuo, Kamyab Javanmardi, Kevin C. Le, et al. 2020. “Structure-Based Design of Prefusion-Stabilized SARS-CoV-2 Spikes.” Science 369 (6510): 1501–5.

Huo, Jiandong, Yuguang Zhao, Jingshan Ren, Daming Zhou, Helen M. E. Duyvesteyn, Helen M. Ginn, Loic Carrique, et al. 2020. “Neutralization of SARS-CoV-2 by Destruction of the Prefusion Spike.” Cell Host & Microbe 28 (3): 445–54.e6.

Juraszek, Jarek, Lucy Rutten, Sven Blokland, Pascale Bouchier, Richard Voorzaat, Tina Ritschel, Mark J. G. Bakkers, Ludovic L. R. Renault, and Johannes P. M. Langedijk. 2021. “Stabilizing the Closed SARS-CoV-2 Spike Trimer.” Nature Communications 12 (1): 244.

Karlsson Hedestam, Gunilla B., Javier Guenaga, Martin Corcoran, and Richard T. Wyatt. 2017. “Evolution of B Cell Analysis and Env Trimer Redesign.” Immunological Reviews 275 (1): 183–202.

Ke, Zunlong, Joaquin Oton, Kun Qu, Mirko Cortese, Vojtech Zila, Lesley McKeane, Takanori Nakane, et al. 2020. “Structures and Distributions of SARS-CoV-2 Spike Proteins on Intact Virions.” Nature 588 (7838): 498–502.

Koenig, Paul-Albert, Hrishikesh Das, Hejun Liu, Beate M. Kümmerer, Florian N. Gohr, Lea-Marie Jenster, Lisa D. J. Schiffelers, et al. 2021. “Structure-Guided Multivalent Nanobodies Block SARS-CoV-2 Infection and Suppress Mutational Escape.” Science 371 (6530). https://doi.org/10.1126/science.abe6230.

Korber, Bette, Will M. Fischer, Sandrasegaram Gnanakaran, Hyejin Yoon, James Theiler, Werner Abfalterer, Nick Hengartner, et al. 2020. “Tracking Changes in SARS-CoV-2 Spike: Evidence That D614G Increases Infectivity of the COVID-19 Virus.” Cell 182, 812–827.

Krammer, Florian. 2020. “SARS-CoV-2 Vaccines in Development.” Nature 586 (7830): 516–27.

Lainšček, Duško, Tina Fink, Vida Forstnerič, Iva Hafner-Bratkovič, Sara Orehek, Žiga Strmšek, Mateja Manček-Keber, et al. 2020. “Immune Response to Vaccine Candidates Based on Different Types of Nanoscaffolded RBD Domain of the SARS-CoV-2 Spike Protein.” *Cold Spring Harbor Laboratory*. https://doi.org/10.1101/2020.08.28.244269.

Liu, Lihong, Pengfei Wang, Manoj S. Nair, Jian Yu, Micah Rapp, Qian Wang, Yang Luo, et al. 2020. “Potent Neutralizing Antibodies against Multiple Epitopes on SARS-CoV-2 Spike.” Nature 584 (7821): 450–56.

Lu, Maolin, Pradeep D. Uchil, Wenwei Li, Desheng Zheng, Daniel S. Terry, Jason Gorman, Wei Shi, et al. 2020. “Real-Time Conformational Dynamics of SARS-CoV-2 Spikes on Virus Particles.” Cell Host & Microbe 28 (6): 880–91.e8.

McLellan, Jason S., Man Chen, M. Gordon Joyce, Mallika Sastry, Guillaume B. E. Stewart-Jones, Yongping Yang, Baoshan Zhang, et al. 2013. “Structure-Based Design of a Fusion Glycoprotein Vaccine for Respiratory Syncytial Virus.” Science 342 (6158): 592–98.

Pallesen, Jesper, Nianshuang Wang, Kizzmekia S. Corbett, Daniel Wrapp, Robert N. Kirchdoerfer, Hannah L. Turner, Christopher A. Cottrell, et al. 2017. “Immunogenicity and Structures of a Rationally Designed Prefusion MERS-CoV Spike Antigen.” Proceedings of the National Academy of Sciences of the United States of America 114 (35): E7348–57.

Planas, Delphine, Timothée Bruel, Ludivine Grzelak, Florence Guivel-Benhassine, Isabelle Staropoli, Françoise Porrot, Cyril Planchais, et al. 2021. “Sensitivity of Infectious SARS-CoV-2 B.1.1.7 and B.1.351 Variants to Neutralizing Antibodies.” Nature Medicine,

Polack, Fernando P., Stephen J. Thomas, Nicholas Kitchin, Judith Absalon, Alejandra Gurtman, Stephen Lockhart, John L. Perez, et al. 2020. “Safety and Efficacy of the BNT162b2 mRNA Covid-19 Vaccine.” The New England Journal of Medicine 383 (27): 2603–15.

Punjani, Ali, John L. Rubinstein, David J. Fleet, and Marcus A. Brubaker. 2017. “cryoSPARC: Algorithms for Rapid Unsupervised Cryo-EM Structure Determination.” Nature Methods 14 (3): 290–96.

Raska, Milan, Kazuo Takahashi, Lydie Czernekova, Katerina Zachova, Stacy Hall, Zina Moldoveanu, Matt C. Elliott, et al. 2010. “Glycosylation Patterns of HIV-1 gp120 Depend on the Type of Expressing Cells and Affect Antibody Recognition.” The Journal of Biological Chemistry 285 (27): 20860–69.

Robbiani, Davide F., Christian Gaebler, Frauke Muecksch, Julio C. C. Lorenzi, Zijun Wang, Alice Cho, Marianna Agudelo, et al. 2020. “Convergent Antibody Responses to SARS-CoV-2 in Convalescent Individuals.” Nature 584 (7821): 437–42.

Rogers, Thomas F., Fangzhu Zhao, Deli Huang, Nathan Beutler, Alison Burns, Wan-Ting He, Oliver Limbo, et al. 2020. “Isolation of Potent SARS-CoV-2 Neutralizing Antibodies and Protection from Disease in a Small Animal Model.” Science 369 (6506): 956–63.

Sacco, Michael Dominic, Chunlong Ma, Panagiotis Lagarias, Ang Gao, Julia Alma Townsend, Xiangzhi Meng, Peter Dube, et al. 2020. “Structure and Inhibition of the SARS-CoV-2 Main Protease Reveal Strategy for Developing Dual Inhibitors against Mpro and Cathepsin L.” Science Advances 6 (50). https://doi.org/10.1126/sciadv.abe0751.

Schoof, Michael, Bryan Faust, Reuben A. Saunders, Smriti Sangwan, Veronica Rezelj, Nick Hoppe, Morgane Boone, et al. 2020. “An Ultrapotent Synthetic Nanobody Neutralizes SARS-CoV-2 by Stabilizing Inactive Spike.” Science 370 (6523): 1473–79.

Shang, Jian, Yushun Wan, Chang Liu, Boyd Yount, Kendra Gully, Yang Yang, Ashley Auerbach, Guiqing Peng, Ralph Baric, and Fang Li. 2020. “Structure of Mouse Coronavirus Spike Protein Complexed with Receptor Reveals Mechanism for Viral Entry.” PLoS Pathogens 16 (3): e1008392.

Shang, Jian, Yushun Wan, Chuming Luo, Gang Ye, Qibin Geng, Ashley Auerbach, and Fang Li. 2020. “Cell Entry Mechanisms of SARS-CoV-2.” Proceedings of the National Academy of Sciences of the United States of America 117 (21): 11727–34.

Sliepen, Kwinten, Thijs van Montfort, Mark Melchers, Gözde Isik, and Rogier W. Sanders. 2015. “Immunosilencing a Highly Immunogenic Protein Trimerization Domain.” The Journal of Biological Chemistry 290 (12): 7436–42.

Stewart-Jones, Guillaume B. E., Paul V. Thomas, Man Chen, Aliaksandr Druz, M. Gordon Joyce, Wing-Pui Kong, Mallika Sastry, et al. 2015. “A Cysteine Zipper Stabilizes a Pre-Fusion F Glycoprotein Vaccine for Respiratory Syncytial Virus.” PloS One 10 (6): e0128779.

Sun, Xiangjie, Akila Jayaraman, Pavithra Maniprasad, Rahul Raman, Katherine V. Houser, Claudia Pappas, Hui Zeng, Ram Sasisekharan, Jacqueline M. Katz, and Terrence M. Tumpey. 2013. “N-Linked Glycosylation of the Hemagglutinin Protein Influences Virulence and Antigenicity of the 1918 Pandemic and Seasonal H1N1 Influenza A Viruses.” Journal of Virology 87 (15): 8756–66.

Sztain, Terra, Surl-Hee Ahn, Anthony T. Bogetti, Lorenzo Casalino, Jory A. Goldsmith, Ryan S. McCool, Fiona L. Kearns, et al. 2021. “A Glycan Gate Controls Opening of the SARS-CoV-2 Spike Protein.” Cold Spring Harbor Laboratory. https://doi.org/10.1101/2021.02.15.431212.

Tegunov, Dimitry, and Patrick Cramer. 2019. “Real-Time Cryo-Electron Microscopy Data Preprocessing with Warp.” Nature Methods 16 (11): 1146–52.

Tortorici, M. Alejandra, Martina Beltramello, Florian A. Lempp, Dora Pinto, Ha V. Dang, Laura E. Rosen, Matthew McCallum, et al. 2020. “Ultrapotent Human Antibodies Protect against SARS-CoV-2 Challenge via Multiple Mechanisms.” Science 370 (6519): 950–57.

Walls, Alexandra C., Young-Jun Park, M. Alejandra Tortorici, Abigail Wall, Andrew T. McGuire, and David Veesler. 2020. “Structure, Function, and Antigenicity of the SARS-CoV-2 Spike Glycoprotein.” Cell 181 (2): 281–92.e6.

Walls, Alexandra C., Xiaoli Xiong, Young-Jun Park, M. Alejandra Tortorici, Joost Snijder, Joel Quispe, Elisabetta Cameroni, et al. 2020. “Unexpected Receptor Functional Mimicry Elucidates Activation of Coronavirus Fusion.” Cell 183 (6): 1732.

Wang, Pengfei, Manoj S. Nair, Lihong Liu, Sho Iketani, Yang Luo, Yicheng Guo, Maple Wang, et al. 2021. “Antibody Resistance of SARS-CoV-2 Variants B.1.351 and B.1.1.7.” Nature, March. https://doi.org/10.1038/s41586-021-03398-2.

Watanabe, Yasunori, Joel D. Allen, Daniel Wrapp, Jason S. McLellan, and Max Crispin. 2020. “Site-Specific Glycan Analysis of the SARS-CoV-2 Spike.” Science 369 (6501): 330–33.

Wrapp, Daniel, Nianshuang Wang, Kizzmekia S. Corbett, Jory A. Goldsmith, Ching-Lin Hsieh, Olubukola Abiona, Barney S. Graham, and Jason S. McLellan. 2020. “Cryo-EM Structure of the 2019-nCoV Spike in the Prefusion Conformation.” Science 367 (6483): 1260–63.

Xiong, Xiaoli, Kun Qu, Katarzyna A. Ciazynska, Myra Hosmillo, Andrew P. Carter, Soraya Ebrahimi, Zunlong Ke, et al. 2020. “A Thermostable, Closed SARS-CoV-2 Spike Protein Trimer.” Nature Structural & Molecular Biology 27 (10): 934–41.

Xu, Cong, Yanxing Wang, Caixuan Liu, Chao Zhang, Wenyu Han, Xiaoyu Hong, Yifan Wang, et al. 2021. “Conformational Dynamics of SARS-CoV-2 Trimeric Spike Glycoprotein in Complex with Receptor ACE2 Revealed by Cryo-EM.” Science Advances 7 (1).

Yao, Hangping, Yutong Song, Yong Chen, Nanping Wu, Jialu Xu, Chujie Sun, Jiaxing Zhang, et al. 2020. “Molecular Architecture of the SARS-CoV-2 Virus.” Cell 183 (3): 730–38.e13.

Zhou, Daming, Wanwisa Dejnirattisai, Piyada Supasa, Chang Liu, Alexander J. Mentzer, Helen M. Ginn, Yuguang Zhao, et al. 2021. “Evidence of Escape of SARS-CoV-2 Variant B.1.351 from Natural and Vaccine-Induced Sera.” Cell,

Zhou, Tongqing, Yaroslav Tsybovsky, Jason Gorman, Micah Rapp, Gabriele Cerutti, Gwo-Yu Chuang, Phinikoula S. Katsamba, et al. 2020. “Cryo-EM Structures of SARS-CoV-2 Spike without and with ACE2 Reveal a pH-Dependent Switch to Mediate Endosomal Positioning of Receptor-Binding Domains.” Cell Host & Microbe 28 (6): 867–79.e5.

