## Supplementary Tables for "Structure-based design of a highly stable, covalently-linked SARS-CoV-2 spike trimer with improved structural properties and immunogenicity"

| Spike variant | Proline backbone | S1/S2 linker | Disulfide bond | Yield (mg/L) | |
|---|---|---|---|---|---|
| | | | | HEK293F | ExpiCHO |
| S-2P | 2P | - | - | 0.7 | 40 |
| S-2P_FL1 | 2P | GGGS | - | 2.3 | ND |
| S-2P_FL2 | 2P | GGGSGGGS | - | 1.9 | 76 |
| HexaPro (6P) | 6P | - | - | 14.5 | 177 |
| 6P_D614G | 6P | - | - | 9.4 | 156 |
| 6P_FL1 | 6P | GGGS | - | 11.6 | 163 |
| 6P_FL2 | 6P | GGGSGGGS | - | 15.6 | 184 |
| 6P_RL3 | 6P | GP | - | 12.1 | 169 |
| 6P_RL4 | 6P | GPGP | - | 8.9 | 172 |
| 6P_DS1 | 6P | - | F42C-G566C | 3 | 110 |
| 6P_DS3 | 6P | - | Y707C-T883C | 4 | 114 |
| 6P_DS5 | 6P | - | A701C-Q787C | 2.3 | 96 |
| 5P | 5P | - | - | 13.2 | 175 |
| 5P_D614G | 5P | - | - | 10.1 | 162 |
| 5P_FL2 | 5P | GGGSGGGS | - | 19.8 | 218 |
| 5P_DS3 | 5P | - | Y707C-T883C | 3.5 | 150 |
| 5P_FL2_DS2 | 5P | GGGSGGGS | G413C-P987C | 2.7 | 86 |
| VFLIP (5P_FL2_DS3) | 5P | GGGSGGGS | Y707C-T883C | 7.1 | 157 |
| VFLIP_D614G | 5P | GGGSGGGS | Y707C-T883C | 8.6 | 128 |

**Extended Data Table S1. Summary of SARS-CoV-2 spike variants used in this study. Related to Figure 1.**

| | RBD | | | | | Hexapro | | | | | VFLIP | | | | | DS2 | | | | |
|---|---|---|---|---|---|---|---|---|---|---|---|---|---|---|---|---|---|---|---|---|
| Name | $k_a$ (M-1 s-1) | $k_d$ (s-1) | $K_D$ (M) | Rmax (RU) | Res SD | $k_a$ (M-1 s-1) | $k_d$ (s-1) | $K_D$ (M) | Rmax (RU) | Res SD | $k_a$ (M-1 s-1) | $k_d$ (s-1) | $K_D$ (M) | Rmax (RU) | Res SD | $k_a$ (M-1 s-1) | $k_d$ (s-1) | $K_D$ (M) | Rmax (RU) | Res SD |
| COVIC-001 | 5.1E+04 | 3.7E-03 | 7.3E-08 | 70.2 | 4.4 | 2.0E+05 | 1.3E-03 | 6.7E-09 | 463.4 | 14.2 | 3.8E+04 | 6.1E-04 | 1.6E-08 | 415.6 | 7.2 | 2.26E+04 | 3.24E-04 | 1.43E-08 | 194.4 | 9.4 |
| COVIC-013 | 7.4E+05 | 3.4E-04 | 4.6E-10 | 91.1 | 3.1 | 3.3E+05 | 4.6E-05 | 1.4E-10 | 358.6 | 11.7 | 7.7E+04 | 1.5E-04 | 1.9E-09 | 164.7 | 7.4 | 5.44E+04 | 8.27E-05 | 1.52E-09 | 90.7 | 7.5 |
| COVIC-015 | 1.7E+05 | 3.1E-03 | 1.9E-08 | 101.7 | 2.0 | 2.1E+05 | 9.0E-05 | 4.3E-10 | 293.0 | 7.6 | 7.4E+04 | 5.1E-05 | 6.9E-10 | 319.4 | 7.8 | 7.19E+04 | 1.57E-04 | 2.19E-09 | 364.3 | 11.0 |
| COVIC-019 | 6.6E+04 | 1.0E-05 | 1.5E-10 | 33.2 | 0.9 | N/A | N/A | N/A | N/A | N/A | N/A | N/A | N/A | N/A | N/A | N/A | N/A | N/A | N/A | N/A |
| COVIC-020 | N/A | N/A | N/A | N/A | N/A | 1.6E+05 | 9.9E-04 | 6.3E-09 | 283.2 | 7.0 | 1.1E+05 | 9.4E-04 | 8.9E-09 | 338.7 | 10.0 | 1.01E+05 | 1.10E-03 | 1.09E-08 | 447.4 | 21.2 |
| COVIC-021 | 1.3E+05 | 1.9E-04 | 1.4E-09 | 72.5 | 2.0 | 9.3E+04 | 4.5E-04 | 4.8E-09 | 66.8 | 2.4 | 2.8E+04 | 1.0E-05 | 3.5E-10 | 65.8 | 2.1 | 1.10E+04 | 1.83E-04 | 1.67E-08 | 75.6 | 2.5 |
| COVIC-022 | 1.3E+05 | 3.6E-02 | 2.7E-07 | 252.2 | 4.5 | 4.8E+04 | 4.5E-04 | 9.4E-09 | 249.7 | 4.6 | 8.2E+03 | 1.8E-04 | 2.3E-08 | 305.3 | 2.0 | 2.79E+04 | 1.97E-03 | 7.07E-08 | 50.2 | 3.4 |
| COVIC-023 | 3.0E+05 | 1.2E-02 | 3.8E-08 | 95.6 | 4.4 | 1.2E+05 | 1.8E-04 | 1.4E-09 | 127.2 | 6.6 | 2.8E+04 | 1.0E-05 | 3.6E-10 | 236.4 | 4.7 | 8.86E+03 | 1.82E-04 | 2.05E-08 | 485.2 | 9.0 |
| COVIC-024 | 3.2E+05 | 1.1E-02 | 3.5E-08 | 165.6 | 4.2 | 4.6E+04 | 4.4E-04 | 9.5E-09 | 230.8 | 2.8 | 1.0E+04 | 1.0E-04 | 1.0E-08 | 286.8 | 2.4 | 1.47E+04 | 4.63E-04 | 3.14E-08 | 62.7 | 1.4 |
| COVIC-028 | 1.9E+05 | 2.5E-03 | 1.3E-08 | 124.2 | 5.8 | 5.5E+04 | 3.1E-04 | 5.6E-09 | 190.3 | 2.0 | 9.9E+03 | 1.8E-04 | 1.8E-08 | 191.8 | 1.5 | N/A | N/A | N/A | N/A | N/A |
| COVIC-037 | N/A | N/A | N/A | N/A | N/A | 1.6E+05 | 1.3E-04 | 8.3E-10 | 38.9 | 1.1 | 8.0E+03 | 1.0E-05 | 1.3E-09 | 50.9 | 3.4 | N/A | N/A | N/A | N/A | N/A |
| COVIC-038 | 1.6E+05 | 4.4E-03 | 2.8E-08 | 60.8 | 2.6 | 9.2E+04 | 3.2E-04 | 3.5E-09 | 137.4 | 2.5 | 2.4E+04 | 2.5E-04 | 1.0E-08 | 99.7 | 1.4 | 7.06E+03 | 1.00E-05 | 1.42E-09 | 47.7 | 7.6 |
| COVIC-043 | 2.0E+05 | 7.8E-03 | 3.9E-08 | 125.5 | 3.3 | 1.5E+05 | 1.8E-04 | 1.2E-09 | 312.4 | 4.8 | 4.1E+04 | 2.4E-04 | 5.9E-09 | 191.1 | 3.2 | 1.40E+04 | 8.06E-05 | 5.76E-09 | 123.1 | 7.1 |
| COVIC-058 | 1.7E+05 | 2.8E-02 | 1.6E-07 | 1121.5 | 18.6 | 1.9E+05 | 6.8E-05 | 3.6E-10 | 1011.5 | 15.4 | 4.7E+04 | 2.3E-05 | 4.9E-10 | 844.7 | 7.0 | 1.55E+04 | 1.54E-04 | 9.92E-09 | 664.6 | 7.3 |
| COVIC-063 | 2.6E+05 | 2.4E-02 | 9.6E-08 | 300.0 | 13.3 | 1.8E+05 | 6.2E-05 | 3.5E-10 | 509.8 | 11.5 | 4.9E+04 | 1.0E-05 | 2.0E-10 | 423.2 | 7.3 | 2.13E+04 | 7.08E-05 | 3.32E-09 | 313.6 | 4.3 |
| COVIC-064 | 1.5E+05 | 1.6E-02 | 1.1E-07 | 83.1 | 4.6 | 1.5E+05 | 2.4E-04 | 1.6E-09 | 235.8 | 2.9 | 3.8E+04 | 3.3E-05 | 8.6E-10 | 207.1 | 2.3 | 1.63E+04 | 1.00E-05 | 6.13E-10 | 92.0 | 6.1 |
| COVIC-065 | 3.7E+05 | 3.2E-02 | 8.7E-08 | 222.5 | 9.2 | 2.1E+05 | 9.3E-05 | 4.5E-10 | 586.1 | 12.5 | 5.8E+04 | 2.8E-05 | 4.8E-10 | 470.8 | 7.2 | 2.59E+04 | 1.34E-05 | 5.19E-10 | 345.0 | 7.4 |
| COVIC-067 | 3.4E+04 | 4.5E-03 | 1.3E-07 | 65.9 | 3.6 | 1.9E+05 | 1.1E-03 | 6.1E-09 | 360.4 | 8.7 | 4.6E+04 | 1.0E-03 | 2.3E-08 | 305.2 | 10.3 | 9.62E+03 | 1.33E-03 | 1.38E-07 | 373.5 | 5.6 |
| COVIC-068 | 1.2E+05 | 8.7E-04 | 7.2E-09 | 59.2 | 1.7 | 1.3E+05 | 8.9E-05 | 6.8E-10 | 181.7 | 2.4 | 5.0E+04 | 1.7E-04 | 3.5E-09 | 111.0 | 2.2 | 4.45E+04 | 2.85E-04 | 6.39E-09 | 110.0 | 2.9 |
| COVIC-073 | 9.6E+04 | 2.3E-03 | 2.4E-08 | 24.0 | 1.0 | 8.3E+04 | 8.3E-04 | 1.0E-08 | 57.3 | 1.4 | 3.8E+03 | 1.1E-03 | 2.9E-07 | 163.8 | 3.3 | 6.90E+03 | 1.10E-03 | 1.60E-07 | 29.6 | 3.7 |
| COVIC-078 | 1.1E+05 | 1.1E-02 | 1.1E-07 | 81.5 | 2.5 | 7.8E+04 | 2.0E-05 | 2.6E-10 | 473.7 | 6.2 | 6.2E+04 | 1.7E-05 | 2.7E-10 | 510.7 | 8.0 | 6.06E+04 | 3.55E-05 | 5.86E-10 | 464.6 | 14.0 |
| COVIC-079 | 1.3E+05 | 3.5E-04 | 2.8E-09 | 154.7 | 2.6 | 1.1E+05 | 1.0E-05 | 8.7E-11 | 406.1 | 3.3 | 3.3E+04 | 1.0E-05 | 3.0E-10 | 371.0 | 6.1 | 8.49E+03 | 1.00E-05 | 1.18E-09 | 583.3 | 2.9 |
| COVIC-082 | 1.8E+05 | 1.9E-02 | 1.1E-07 | 200.2 | 3.3 | 3.0E+04 | 5.0E-04 | 1.7E-08 | 243.2 | 3.7 | 6.0E+03 | 3.3E-04 | 5.5E-08 | 341.3 | 2.6 | 7.00E+04 | 4.94E-03 | 7.05E-08 | 72.8 | 7.6 |
| COVIC-091 | 1.4E+05 | 2.2E-03 | 1.5E-08 | 148.5 | 2.4 | 4.0E+04 | 3.0E-04 | 7.5E-09 | 374.7 | 3.8 | 3.5E+04 | 1.0E-05 | 2.9E-10 | 231.5 | 5.4 | 1.26E+04 | 1.00E-05 | 7.93E-10 | 208.5 | 13.7 |
| COVIC-094 | 5.9E+04 | 7.0E-04 | 1.2E-08 | 508.0 | 6.7 | 7.8E+04 | 1.0E-05 | 1.3E-10 | 547.9 | 3.8 | 2.8E+04 | 1.0E-05 | 3.6E-10 | 515.4 | 5.8 | 1.82E+04 | 1.00E-05 | 5.48E-10 | 354.4 | 19.8 |
| COVIC-094 | 7.3E+04 | 7.2E-04 | 9.9E-09 | 168.6 | 2.4 | 4.8E+04 | 1.0E-05 | 2.1E-10 | 368.0 | 4.2 | 1.2E+04 | 7.8E-05 | 6.6E-09 | 483.0 | 5.4 | 3.78E+03 | 9.80E-05 | 2.60E-08 | 699.1 | 6.5 |

| | | | | | | | | | | | | | | | | | | | | |
|---|---|---|---|---|---|---|---|---|---|---|---|---|---|---|---|---|---|---|---|---|
| **COVIC-095** | 8.5E+04 | 5.7E-03 | 6.6E-08 | 427.2 | 5.2 | 9.2E+04 | 6.9E-05 | 7.5E-10 | 613.5 | 2.8 | 3.5E+04 | 1.0E-05 | 2.9E-10 | 584.6 | 8.6 | 1.81E+04 | 1.00E-05 | 5.53E-10 | 525.7 | 14.6 |
| **COVIC-096** | 7.1E+04 | 1.1E-02 | 1.5E-07 | 486.3 | 3.5 | 1.7E+05 | 3.9E-05 | 2.3E-10 | 789.3 | 15.1 | 9.4E+04 | 1.0E-05 | 1.1E-10 | 952.2 | 35.1 | 8.61E+04 | 1.00E-05 | 1.16E-10 | 911.0 | 52.0 |
| **COVIC-096** | 8.3E+04 | 1.4E-02 | 1.6E-07 | 135.4 | 1.7 | 1.4E+05 | 9.7E-05 | 7.0E-10 | 400.6 | 5.7 | 8.1E+04 | 2.9E-05 | 3.5E-10 | 453.1 | 12.4 | 7.03E+04 | 1.00E-05 | 1.42E-10 | 421.6 | 18.6 |
| **COVIC-097** | 4.6E+05 | 4.0E-03 | 8.6E-09 | 502.3 | 11.4 | 3.6E+05 | 1.0E-04 | 2.8E-10 | 951.1 | 35.6 | 1.1E+05 | 2.3E-05 | 2.2E-10 | 690.4 | 27.0 | 9.21E+04 | 7.16E-05 | 7.77E-10 | 601.0 | 26.2 |
| **COVIC-097** | 2.2E+05 | 2.1E-03 | 9.2E-09 | 136.1 | 1.6 | 1.2E+05 | 1.9E-04 | 1.6E-09 | 402.1 | 3.8 | 2.6E+04 | 1.3E-04 | 4.9E-09 | 450.5 | 6.0 | 1.82E+03 | 5.89E-04 | 3.23E-07 | 2110.0 | 7.5 |
| **COVIC-098** | 1.3E+05 | 1.6E-03 | 1.2E-08 | 510.3 | 8.6 | 1.6E+05 | 8.4E-05 | 5.3E-10 | 668.7 | 8.3 | 4.5E+04 | 6.9E-05 | 1.5E-09 | 683.3 | 7.4 | 1.32E+04 | 3.24E-04 | 2.45E-08 | 723.3 | 5.6 |
| **COVIC-099** | 1.5E+05 | 3.5E-04 | 2.3E-09 | 409.9 | 10.2 | 2.0E+05 | 4.5E-05 | 2.2E-10 | 724.6 | 22.6 | 6.3E+04 | 2.3E-05 | 3.8E-10 | 472.5 | 10.5 | 5.42E+04 | 1.00E-05 | 1.84E-10 | 391.8 | 18.9 |
| **COVIC-100** | 4.0E+05 | 3.2E-03 | 8.0E-09 | 521.4 | 13.5 | 4.1E+05 | 1.0E-04 | 2.5E-10 | 1012.9 | 42.1 | 1.6E+05 | 1.9E-05 | 1.2E-10 | 926.5 | 46.8 | 1.14E+05 | 1.80E-05 | 1.59E-10 | 936.6 | 41.5 |
| **COVIC-101** | 8.8E+04 | 8.2E-04 | 9.4E-09 | 534.7 | 6.2 | 2.1E+05 | 1.2E-05 | 5.8E-11 | 904.1 | 19.2 | 1.1E+05 | 1.0E-05 | 9.0E-11 | 976.9 | 36.7 | 1.29E+05 | 1.00E-05 | 7.73E-11 | 1024.0 | 59.4 |
| **COVIC-102** | 1.1E+05 | 4.8E-04 | 4.2E-09 | 576.0 | 21.0 | 2.0E+05 | 1.0E-05 | 5.0E-11 | 259.2 | 17.7 | 2.9E+04 | 1.0E-05 | 3.5E-10 | 195.1 | 7.7 | 1.00E+04 | 1.00E-04 | 1.00E-08 | 0.0 | 50.7 |
| **COVIC-103** | 1.6E+05 | 4.5E-04 | 2.9E-09 | 588.4 | 14.2 | 1.8E+05 | 1.0E-05 | 5.5E-11 | 332.7 | 22.2 | 4.8E+04 | 1.0E-05 | 2.1E-10 | 145.9 | 11.7 | 1.00E+04 | 1.00E-04 | 1.00E-08 | 0.0 | 43.5 |
| **COVIC-104** | 1.6E+05 | 1.6E-04 | 1.0E-09 | 98.2 | 8.4 | 1.5E+05 | 9.5E-05 | 6.3E-10 | 876.1 | 14.0 | 7.9E+04 | 1.4E-04 | 1.8E-09 | 596.1 | 20.4 | 9.65E+04 | 7.29E-05 | 7.55E-10 | 488.3 | 29.1 |
| **COVIC-105** | 3.2E+05 | 6.1E-03 | 1.9E-08 | 494.5 | 9.5 | 3.2E+05 | 1.1E-04 | 3.4E-10 | 950.3 | 33.5 | 1.1E+05 | 1.0E-05 | 8.7E-11 | 861.9 | 38.6 | 1.09E+05 | 1.00E-05 | 9.19E-11 | 885.9 | 47.3 |
| **COVIC-106** | 4.0E+05 | 1.1E-03 | 2.8E-09 | 437.7 | 10.3 | 3.1E+05 | 6.7E-05 | 2.1E-10 | 751.7 | 30.5 | 7.7E+04 | 2.0E-05 | 2.6E-10 | 472.8 | 17.4 | 9.29E+04 | 1.00E-05 | 1.08E-10 | 368.3 | 21.8 |
| **COVIC-107** | 2.9E+05 | 9.2E-04 | 3.2E-09 | 465.7 | 11.0 | 2.6E+05 | 3.5E-05 | 1.3E-10 | 726.2 | 22.2 | 6.6E+04 | 1.0E-05 | 1.5E-10 | 470.0 | 13.6 | 6.44E+04 | 1.00E-05 | 1.55E-10 | 330.7 | 31.9 |
| **COVIC-108** | 3.3E+05 | 9.5E-04 | 2.9E-09 | 494.6 | 11.5 | 2.8E+05 | 7.4E-05 | 2.7E-10 | 848.8 | 28.5 | 8.3E+04 | 1.0E-05 | 1.2E-10 | 610.1 | 21.8 | 8.84E+04 | 1.00E-05 | 1.13E-10 | 510.7 | 49.7 |
| **COVIC-109** | 2.8E+05 | 4.4E-04 | 1.5E-09 | 538.6 | 14.1 | 3.3E+05 | 2.5E-05 | 7.4E-11 | 891.8 | 29.8 | 1.1E+05 | 1.0E-05 | 9.1E-11 | 654.0 | 23.7 | 1.01E+05 | 1.00E-05 | 9.89E-11 | 600.8 | 45.5 |
| **COVIC-110** | 2.8E+05 | 1.4E-03 | 5.1E-09 | 472.1 | 8.9 | 3.1E+05 | 8.8E-05 | 2.8E-10 | 879.3 | 31.3 | 1.2E+05 | 1.0E-05 | 8.5E-11 | 768.0 | 32.5 | 1.12E+05 | 1.00E-05 | 8.91E-11 | 767.4 | 52.8 |
| **COVIC-111** | 2.6E+05 | 8.4E-04 | 3.2E-09 | 537.1 | 14.6 | 3.4E+05 | 4.1E-05 | 1.2E-10 | 949.2 | 34.8 | 1.3E+05 | 1.0E-05 | 7.5E-11 | 892.5 | 39.6 | 1.21E+05 | 1.00E-05 | 8.26E-11 | 888.2 | 66.9 |
| **COVIC-112** | 2.6E+05 | 5.0E-04 | 1.9E-09 | 513.1 | 16.8 | 3.0E+05 | 1.0E-05 | 3.3E-11 | 795.4 | 27.0 | 8.0E+04 | 1.5E-05 | 1.9E-10 | 567.2 | 21.1 | 7.43E+04 | 1.00E-05 | 1.35E-10 | 449.9 | 33.6 |
| **COVIC-113** | 1.1E+05 | 1.2E-03 | 1.1E-08 | 349.8 | 13.7 | 5.2E+04 | 1.8E-04 | 3.4E-09 | 483.7 | 6.7 | 1.2E+04 | 6.3E-05 | 5.3E-09 | 583.0 | 5.4 | 1.59E+03 | 2.16E-03 | 1.36E-06 | 1009.8 | 8.3 |
| **COVIC-114** | 1.3E+05 | 6.9E-04 | 5.3E-09 | 368.1 | 13.5 | 6.3E+04 | 4.0E-05 | 6.3E-10 | 386.5 | 3.0 | 1.4E+04 | 1.0E-05 | 7.1E-10 | 462.1 | 4.9 | 5.48E+03 | 1.00E-05 | 1.82E-09 | 149.1 | 14.0 |
| **COVIC-115** | 1.6E+05 | 5.3E-04 | 3.4E-09 | 296.9 | 11.0 | 6.4E+04 | 1.2E-04 | 1.9E-09 | 343.9 | 2.0 | 1.6E+04 | 2.1E-05 | 1.3E-09 | 389.3 | 2.2 | 1.18E+04 | 3.05E-04 | 2.59E-08 | 99.5 | 2.0 |
| **COVIC-116** | 5.1E+04 | 1.1E-03 | 2.2E-08 | 443.1 | 3.5 | 1.3E+05 | 5.9E-05 | 4.4E-10 | 671.7 | 10.0 | 7.1E+04 | 1.0E-05 | 1.4E-10 | 576.2 | 19.0 | 5.29E+04 | 1.00E-05 | 1.89E-10 | 586.3 | 38.2 |
| **COVIC-117** | 3.7E+05 | 1.6E-03 | 4.4E-09 | 468.6 | 8.7 | 4.1E+05 | 9.0E-05 | 2.2E-10 | 988.5 | 34.0 | 2.1E+05 | 1.0E-05 | 4.8E-11 | 979.0 | 51.2 | 2.07E+05 | 1.00E-05 | 4.84E-11 | 1101.9 | 65.9 |
| **COVIC-118** | 3.8E+05 | 8.0E-04 | 2.1E-09 | 179.4 | 13.2 | 3.0E+05 | 7.4E-05 | 2.4E-10 | 586.3 | 16.3 | 1.1E+05 | 6.8E-05 | 6.0E-10 | 462.9 | 15.1 | 4.93E+04 | 4.42E-05 | 8.97E-10 | 405.3 | 11.1 |
| **COVIC-119** | 1.8E+05 | 2.2E-02 | 1.2E-07 | 1225.5 | 20.2 | 1.8E+05 | 5.7E-05 | 3.2E-10 | 1026.5 | 17.5 | 5.1E+04 | 3.0E-05 | 5.8E-10 | 799.1 | 9.6 | 2.41E+04 | 9.40E-05 | 3.91E-09 | 509.3 | 5.2 |
| **COVIC-121** | 1.7E+05 | 2.0E-02 | 1.2E-07 | 849.7 | 18.2 | 1.5E+05 | 8.6E-05 | 5.9E-10 | 777.2 | 12.9 | 3.1E+04 | 2.1E-05 | 6.8E-10 | 596.6 | 6.4 | 1.30E+04 | 1.00E-05 | 7.67E-10 | 343.1 | 7.1 |

| | | | | | | | | | | | | | | | | | | | | |
|---|---|---|---|---|---|---|---|---|---|---|---|---|---|---|---|---|---|---|---|---|
| **COVIC-122** | 1.4E+05 | 1.8E-02 | 1.3E-07 | 1341.9 | 21.6 | 1.8E+05 | 7.9E-05 | 4.4E-10 | 872.7 | 17.0 | 4.4E+04 | 3.0E-05 | 6.7E-10 | 659.2 | 7.4 | 2.06E+04 | 9.16E-05 | 4.45E-09 | 364.3 | 5.5 |
| **COVIC-123** | 1.4E+05 | 1.8E-02 | 1.2E-07 | 1813.3 | 22.7 | 2.0E+05 | 8.6E-05 | 4.4E-10 | 1111.8 | 21.5 | 5.3E+04 | 1.0E-05 | 1.9E-10 | 887.2 | 12.5 | 2.31E+04 | 8.43E-05 | 3.64E-09 | 551.6 | 8.5 |
| **COVIC-124** | 1.2E+05 | 1.5E-02 | 1.2E-07 | 2103.4 | 24.3 | 1.9E+05 | 7.6E-05 | 3.9E-10 | 1186.5 | 19.3 | 5.6E+04 | 2.5E-05 | 4.4E-10 | 984.0 | 11.4 | 2.65E+04 | 3.27E-05 | 1.23E-09 | 662.2 | 10.0 |
| **COVIC-125** | 1.2E+05 | 1.8E-02 | 1.5E-07 | 1882.4 | 22.1 | 2.0E+05 | 6.5E-05 | 3.3E-10 | 1111.8 | 23.7 | 5.5E+04 | 2.3E-05 | 4.2E-10 | 953.1 | 14.7 | 3.06E+04 | 4.65E-05 | 1.52E-09 | 571.9 | 17.3 |
| **COVIC-126** | 1.5E+05 | 2.2E-02 | 1.4E-07 | 1616.1 | 18.5 | 2.3E+05 | 8.5E-05 | 3.8E-10 | 1112.5 | 25.1 | 6.5E+04 | 1.0E-05 | 1.6E-10 | 963.1 | 15.1 | 3.44E+04 | 9.59E-05 | 2.78E-09 | 660.4 | 8.2 |
| **COVIC-127** | 1.1E+05 | 1.4E-02 | 1.3E-07 | 2386.7 | 24.3 | 2.0E+05 | 8.6E-05 | 4.3E-10 | 1270.3 | 27.0 | 5.3E+04 | 2.8E-05 | 5.3E-10 | 1018.5 | 13.9 | 3.00E+04 | 1.69E-04 | 5.62E-09 | 610.1 | 9.1 |
| **COVIC-128** | 1.9E+05 | 2.3E-02 | 1.2E-07 | 1206.8 | 18.1 | 1.8E+05 | 3.8E-05 | 2.1E-10 | 1138.6 | 18.3 | 5.1E+04 | 2.1E-05 | 4.1E-10 | 979.8 | 11.7 | 2.53E+04 | 8.63E-05 | 3.41E-09 | 643.1 | 9.4 |
| **COVIC-129** | 1.4E+05 | 1.4E-02 | 1.1E-07 | 2075.2 | 25.4 | 2.1E+05 | 7.2E-05 | 3.4E-10 | 1328.9 | 28.2 | 5.7E+04 | 2.1E-05 | 3.7E-10 | 1084.9 | 15.0 | 3.10E+04 | 1.22E-04 | 3.94E-09 | 674.1 | 8.3 |
| **COVIC-131** | 1.1E+05 | 1.0E-03 | 8.9E-09 | 393.2 | 6.0 | 3.1E+04 | 1.1E-04 | 3.7E-09 | 270.9 | 2.0 | 2.1E+03 | 1.0E-05 | 4.8E-09 | 996.5 | 4.0 | 9.92E+03 | 6.89E-04 | 6.95E-08 | 65.7 | 2.4 |
| **COVIC-132** | N/A | N/A | N/A | N/A | N/A | 7.6E+04 | 1.6E-04 | 2.1E-09 | 465.3 | 7.4 | 8.9E+03 | 1.3E-04 | 1.4E-08 | 668.1 | 4.8 | 2.52E+04 | 1.00E-05 | 3.97E-10 | 425.6 | 15.1 |
| **COVIC-133** | N/A | N/A | N/A | N/A | N/A | 3.1E+04 | 4.7E-04 | 1.5E-08 | 297.7 | 2.2 | 6.7E+03 | 2.7E-04 | 4.0E-08 | 548.7 | 3.9 | 2.19E+04 | 3.75E-04 | 1.71E-08 | 126.1 | 4.1 |
| **COVIC-134** | 2.1E+04 | 6.2E-03 | 2.9E-07 | 646.1 | 4.6 | 1.2E+05 | 5.4E-05 | 4.5E-10 | 642.7 | 10.8 | 5.2E+04 | 1.0E-05 | 1.9E-10 | 793.1 | 21.5 | 6.39E+04 | 1.00E-05 | 1.56E-10 | 750.6 | 53.5 |
| **COVIC-135** | N/A | N/A | N/A | N/A | N/A | N/A | N/A | N/A | N/A | N/A | N/A | N/A | N/A | N/A | N/A | N/A | N/A | N/A | N/A | N/A |
| **COVIC-136** | 9.6E+04 | 1.8E-03 | 1.8E-08 | 229.2 | 1.9 | 1.1E+05 | 6.9E-05 | 6.4E-10 | 463.1 | 4.5 | 3.0E+04 | 1.0E-05 | 3.3E-10 | 467.2 | 4.8 | 1.27E+04 | 2.06E-04 | 1.63E-08 | 437.9 | 6.9 |
| **COVIC-137** | N/A | N/A | N/A | N/A | N/A | 4.0E+04 | 5.5E-05 | 1.4E-09 | 547.9 | 4.0 | 1.0E+04 | 1.0E-05 | 9.8E-10 | 408.5 | 3.9 | 4.33E+04 | 1.22E-03 | 2.82E-08 | 63.2 | 7.0 |
| **COVIC-138** | N/A | N/A | N/A | N/A | N/A | 2.0E+04 | 1.7E-04 | 8.3E-09 | 323.0 | 2.5 | 1.2E+03 | 1.4E-04 | 1.2E-07 | 1634.1 | 2.7 | 1.22E+04 | 2.21E-04 | 1.81E-08 | 209.7 | 4.1 |
| **COVIC-139** | 1.2E+05 | 1.7E-02 | 1.5E-07 | 455.8 | 8.4 | 3.4E+04 | 1.6E-04 | 4.7E-09 | 537.3 | 6.3 | 6.0E+03 | 2.2E-04 | 3.7E-08 | 772.5 | 4.1 | 1.86E+04 | 1.36E-03 | 7.30E-08 | 111.1 | 4.8 |
| **COVIC-140** | 3.1E+04 | 7.2E-03 | 2.3E-07 | 368.4 | 15.3 | 3.1E+05 | 4.5E-05 | 1.4E-10 | 1148.8 | 36.0 | 1.7E+05 | 4.2E-05 | 2.5E-10 | 1226.1 | 59.9 | 1.44E+05 | 5.33E-05 | 3.70E-10 | 1187.5 | 59.1 |
| **COVIC-141** | 2.6E+05 | 2.6E-02 | 1.0E-07 | 404.7 | 8.7 | 2.4E+05 | 1.9E-05 | 7.9E-11 | 987.2 | 27.6 | 9.9E+04 | 1.0E-05 | 1.0E-10 | 1006.4 | 36.4 | 8.52E+04 | 1.00E-05 | 1.17E-10 | 916.0 | 46.7 |
| **COVIC-142** | 1.1E+05 | 2.4E-03 | 2.1E-08 | 363.6 | 6.3 | 1.2E+05 | 9.4E-05 | 7.7E-10 | 681.3 | 8.2 | 3.6E+04 | 1.0E-05 | 2.8E-10 | 639.6 | 6.0 | 1.38E+04 | 9.39E-05 | 6.80E-09 | 553.0 | 13.4 |
| **COVIC-143** | 1.4E+05 | 2.6E-03 | 1.8E-08 | 496.9 | 5.2 | 1.8E+05 | 1.0E-04 | 5.5E-10 | 758.4 | 12.7 | 5.6E+04 | 1.9E-05 | 3.4E-10 | 751.2 | 10.7 | 2.02E+04 | 4.11E-05 | 2.03E-09 | 659.9 | 10.7 |
| **COVIC-144** | 4.8E+04 | 7.2E-05 | 1.5E-09 | 318.4 | 7.5 | 6.3E+04 | 2.2E-05 | 3.5E-10 | 531.3 | 3.3 | 3.9E+04 | 1.0E-05 | 2.6E-10 | 425.9 | 11.1 | 2.63E+04 | 9.18E-05 | 3.49E-09 | 350.9 | 4.9 |
| **COVIC-145** | 2.3E+05 | 2.7E-02 | 1.2E-07 | 746.2 | 14.6 | 1.7E+05 | 8.7E-05 | 5.1E-10 | 944.6 | 20.2 | 4.7E+04 | 1.0E-05 | 2.1E-10 | 752.8 | 11.2 | 2.82E+04 | 1.64E-05 | 5.82E-10 | 419.4 | 10.6 |
| **COVIC-146** | N/A | N/A | N/A | N/A | N/A | 2.0E+04 | 1.0E-05 | 4.9E-10 | 418.7 | 2.2 | 2.8E+01 | 1.0E-05 | 3.6E-07 | 61647.4 | 4.4 | 5.57E+04 | 1.61E-03 | 2.90E-08 | 30.8 | 3.3 |
| **COVIC-147** | 3.8E+05 | 1.8E-03 | 4.7E-09 | 579.7 | 13.8 | 5.1E+05 | 6.3E-05 | 1.2E-10 | 1126.8 | 51.0 | 2.3E+05 | 1.0E-05 | 4.4E-11 | 1086.8 | 61.2 | 1.49E+05 | 1.11E-04 | 7.47E-10 | 1073.2 | 52.7 |
| **COVIC-148** | 2.5E+05 | 1.3E-03 | 5.0E-09 | 516.9 | 7.9 | 3.2E+05 | 6.0E-05 | 1.9E-10 | 958.8 | 31.6 | 1.2E+05 | 1.0E-05 | 8.1E-11 | 895.2 | 35.4 | 1.02E+05 | 1.00E-05 | 9.84E-11 | 825.3 | 54.6 |
| **COVIC-149** | 3.0E+05 | 1.3E-02 | 4.3E-08 | 535.6 | 11.3 | 4.0E+05 | 2.5E-05 | 6.3E-11 | 1156.1 | 42.1 | 3.0E+05 | 1.0E-05 | 3.4E-11 | 1268.9 | 77.2 | 2.80E+05 | 1.00E-05 | 3.57E-11 | 1290.2 | 91.3 |
| **COVIC-150** | 7.1E+04 | 2.2E-02 | 3.0E-07 | 548.7 | 8.3 | 1.4E+05 | 7.2E-05 | 5.3E-10 | 735.5 | 10.8 | 5.0E+04 | 2.6E-04 | 5.3E-09 | 698.7 | 7.2 | 2.47E+04 | 1.76E-04 | 7.12E-09 | 458.5 | 16.2 |

| | | | | | | | | | | | | | | | | | | | | |
|---|---|---|---|---|---|---|---|---|---|---|---|---|---|---|---|---|---|---|---|---|
| **COVIC-151** | 2.0E+05 | 4.2E-03 | 2.1E-08 | 513.0 | 9.1 | 8.7E+04 | 4.7E-04 | 5.3E-09 | 569.5 | 7.3 | 3.8E+04 | 1.1E-04 | 3.0E-09 | 626.1 | 7.0 | 2.28E+04 | 1.00E-05 | 4.38E-10 | 612.5 | 11.0 |
| **COVIC-152** | 1.6E+05 | 1.4E-03 | 8.6E-09 | 526.4 | 12.8 | 8.1E+04 | 2.6E-04 | 3.2E-09 | 514.9 | 4.0 | 3.0E+04 | 1.2E-04 | 4.1E-09 | 649.1 | 4.6 | 2.02E+04 | 1.43E-04 | 7.05E-09 | 671.9 | 9.7 |
| **COVIC-153** | 1.1E+05 | 2.2E-04 | 2.0E-09 | 443.4 | 11.8 | 1.5E+05 | 3.6E-05 | 2.4E-10 | 678.8 | 10.7 | 4.7E+04 | 1.0E-05 | 2.1E-10 | 698.2 | 9.8 | 2.06E+04 | 1.75E-04 | 8.49E-09 | 678.5 | 9.6 |
| **COVIC-154** | N/A | N/A | N/A | N/A | N/A | 9.0E+04 | 2.6E-04 | 3.0E-09 | 64.1 | 1.4 | N/A | N/A | N/A | N/A | N/A | 3.07E+04 | 1.00E-03 | 3.27E-08 | 40.4 | 2.8 |
| **COVIC-155** | 1.5E+05 | 6.0E-04 | 4.1E-09 | 216.8 | 7.2 | 1.4E+05 | 1.0E-05 | 7.1E-11 | 188.1 | 9.2 | 2.0E+04 | 4.2E-05 | 2.0E-09 | 89.4 | 2.0 | 1.48E+04 | 9.15E-04 | 6.17E-08 | 58.9 | 3.0 |
| **COVIC-156** | 1.3E+05 | 1.8E-04 | 1.4E-09 | 552.1 | 14.2 | 1.9E+05 | 1.0E-05 | 5.3E-11 | 794.9 | 12.7 | 6.1E+04 | 1.5E-05 | 2.5E-10 | 697.6 | 10.8 | 1.94E+04 | 1.00E-05 | 5.15E-10 | 771.2 | 12.8 |
| **COVIC-157** | 8.3E+04 | 1.0E-03 | 1.3E-08 | 498.7 | 7.5 | 1.3E+05 | 1.0E-04 | 7.5E-10 | 669.6 | 11.0 | 4.1E+04 | 2.0E-05 | 4.9E-10 | 664.8 | 7.9 | 1.53E+04 | 4.52E-05 | 2.95E-09 | 661.0 | 13.7 |
| **COVIC-158** | 1.1E+05 | 2.5E-04 | 2.3E-09 | 444.3 | 11.7 | 1.3E+05 | 1.0E-05 | 7.5E-11 | 615.1 | 13.8 | 2.8E+04 | 3.8E-05 | 1.4E-09 | 451.7 | 6.3 | 3.23E+03 | 6.53E-05 | 2.02E-08 | 1259.9 | 7.1 |
| **COVIC-159** | 1.3E+05 | 6.2E-03 | 4.7E-08 | 530.0 | 8.0 | 1.8E+05 | 9.0E-05 | 5.0E-10 | 792.6 | 14.5 | 5.7E+04 | 2.5E-05 | 4.3E-10 | 812.7 | 15.7 | 2.10E+04 | 2.01E-04 | 9.56E-09 | 728.0 | 11.1 |
| **COVIC-160** | 7.9E+04 | 2.6E-03 | 3.3E-08 | 546.4 | 3.8 | 1.4E+05 | 1.3E-04 | 8.8E-10 | 715.8 | 9.4 | 4.2E+04 | 4.8E-05 | 1.1E-09 | 728.9 | 7.7 | 1.62E+04 | 3.20E-04 | 1.97E-08 | 741.6 | 14.2 |
| **COVIC-161** | N/A | N/A | N/A | N/A | N/A | 1.4E+04 | 2.5E-04 | 1.8E-08 | 439.2 | 1.6 | 3.7E+03 | 1.9E-04 | 5.1E-08 | 908.2 | 4.9 | 1.06E+04 | 4.29E-04 | 4.03E-08 | 537.6 | 8.7 |
| **COVIC-162** | 4.6E+04 | 2.8E-02 | 6.2E-07 | 405.4 | 5.3 | 4.2E+04 | 2.4E-04 | 5.7E-09 | 519.3 | 4.1 | 7.3E+03 | 3.1E-04 | 4.2E-08 | 638.1 | 4.4 | 8.29E+03 | 1.26E-03 | 1.52E-07 | 190.2 | 4.1 |
| **COVIC-163** | 6.4E+04 | 1.9E-02 | 2.9E-07 | 486.6 | 4.8 | 1.1E+05 | 4.4E-05 | 3.9E-10 | 654.3 | 9.2 | 3.7E+04 | 1.0E-04 | 2.7E-09 | 673.2 | 5.5 | 1.28E+04 | 2.88E-04 | 2.25E-08 | 576.9 | 8.0 |
| **COVIC-164** | 8.2E+04 | 4.4E-04 | 5.4E-09 | 518.4 | 5.9 | 1.2E+05 | 3.2E-05 | 2.6E-10 | 699.0 | 6.5 | 3.9E+04 | 1.0E-05 | 2.6E-10 | 715.9 | 7.8 | 1.46E+04 | 1.00E-04 | 6.84E-09 | 779.5 | 8.9 |
| **COVIC-165** | N/A | N/A | N/A | N/A | N/A | 7.6E+04 | 1.6E-03 | 2.2E-08 | 68.5 | 3.3 | 3.5E+04 | 3.8E-03 | 1.1E-07 | 159.1 | 7.6 | 9.47E+04 | 6.98E-04 | 7.37E-09 | 100.8 | 9.5 |
| **COVIC-166** | 2.6E+05 | 7.7E-04 | 2.9E-09 | 428.0 | 22.4 | 2.5E+05 | 9.8E-05 | 3.9E-10 | 567.2 | 14.5 | 7.2E+04 | 1.1E-05 | 1.5E-10 | 636.7 | 10.5 | 3.78E+04 | 5.60E-05 | 1.48E-09 | 675.4 | 9.6 |
| **COVIC-167** | 9.9E+04 | 2.4E-04 | 2.5E-09 | 465.4 | 11.1 | 1.6E+05 | 1.3E-05 | 8.1E-11 | 663.2 | 10.3 | 4.8E+04 | 1.0E-05 | 2.1E-10 | 684.9 | 12.9 | 1.80E+04 | 1.00E-05 | 5.55E-10 | 678.0 | 5.4 |
| **COVIC-168** | 1.4E+05 | 5.2E-03 | 3.6E-08 | 459.9 | 10.2 | 2.0E+05 | 9.0E-05 | 4.6E-10 | 791.8 | 16.7 | 6.2E+04 | 1.0E-05 | 1.6E-10 | 705.1 | 15.9 | 2.51E+04 | 2.48E-04 | 9.86E-09 | 642.7 | 11.5 |
| **COVIC-169** | 1.9E+05 | 9.8E-03 | 5.1E-08 | 465.1 | 5.4 | 1.8E+05 | 5.0E-05 | 2.8E-10 | 851.0 | 18.3 | 6.4E+04 | 1.0E-05 | 1.6E-10 | 818.7 | 18.9 | 2.56E+04 | 2.90E-04 | 1.13E-08 | 758.8 | 13.9 |
| **COVIC-170** | 1.4E+05 | 7.5E-04 | 5.4E-09 | 481.5 | 10.5 | 7.1E+04 | 1.4E-04 | 1.9E-09 | 440.5 | 2.7 | 2.6E+04 | 2.7E-05 | 1.1E-09 | 559.4 | 2.2 | 1.15E+04 | 1.00E-05 | 8.66E-10 | 687.7 | 4.9 |
| **COVIC-171** | N/A | N/A | N/A | N/A | N/A | 1.9E+05 | 4.4E-05 | 2.4E-10 | 734.7 | 12.8 | 3.0E+04 | 6.5E-05 | 2.2E-09 | 649.0 | 5.5 | 1.01E+04 | 8.91E-04 | 8.85E-08 | 82.9 | 2.4 |
| **COVIC-172** | 1.3E+05 | 3.1E-03 | 2.4E-08 | 478.1 | 5.6 | 5.4E+04 | 4.1E-04 | 7.6E-09 | 542.4 | 4.4 | 2.6E+04 | 1.1E-04 | 4.3E-09 | 591.4 | 6.2 | 1.80E+04 | 1.00E-05 | 5.56E-10 | 550.0 | 9.5 |
| **COVIC-173** | N/A | N/A | N/A | N/A | N/A | 8.2E+04 | 4.6E-05 | 5.6E-10 | 730.9 | 7.8 | 5.0E+04 | 1.0E-05 | 2.0E-10 | 851.3 | 12.5 | 6.02E+04 | 1.00E-05 | 1.66E-10 | 870.6 | 32.5 |
| **COVIC-174** | N/A | N/A | N/A | N/A | N/A | N/A | N/A | N/A | N/A | N/A | N/A | N/A | N/A | N/A | N/A | N/A | N/A | N/A | N/A | N/A |
| **COVIC-175** | N/A | N/A | N/A | N/A | N/A | N/A | N/A | N/A | N/A | N/A | N/A | N/A | N/A | N/A | N/A | N/A | N/A | N/A | N/A | N/A |
| **COVIC-176** | N/A | N/A | N/A | N/A | N/A | N/A | N/A | N/A | N/A | N/A | N/A | N/A | N/A | N/A | N/A | N/A | N/A | N/A | N/A | N/A |
| **COVIC-177** | N/A | N/A | N/A | N/A | N/A | 1.8E+05 | 1.1E-03 | 6.1E-09 | 18.1 | 1.6 | N/A | N/A | N/A | N/A | N/A | N/A | N/A | N/A | N/A | N/A |
| **COVIC-178** | 1.7E+05 | 1.6E-02 | 9.6E-08 | 311.8 | 3.0 | 7.8E+04 | 3.0E-04 | 3.9E-09 | 296.1 | 3.9 | 2.9E+04 | 3.5E-05 | 1.2E-09 | 338.4 | 4.8 | 1.52E+04 | 1.00E-05 | 6.57E-10 | 391.9 | 9.1 |

| | | | | | | | | | | | | | | | | | | | | |
|---|---|---|---|---|---|---|---|---|---|---|---|---|---|---|---|---|---|---|---|---|
| **COVIC-179** | 1.1E+05 | 4.7E-03 | 4.2E-08 | 366.9 | 3.8 | 1.4E+05 | 4.1E-05 | 2.9E-10 | 704.7 | 8.2 | 5.6E+04 | 1.0E-05 | 1.8E-10 | 713.0 | 13.0 | 4.07E+04 | 1.00E-05 | 2.46E-10 | 666.6 | 21.6 |
| **COVIC-180** | 2.0E+04 | 4.5E-03 | 2.3E-07 | 124.0 | 4.4 | 2.9E+05 | 3.1E-04 | 1.1E-09 | 885.9 | 17.3 | 2.1E+05 | 9.9E-04 | 4.8E-09 | 899.1 | 40.0 | 1.67E+05 | 4.35E-04 | 2.61E-09 | 763.7 | 43.5 |
| **COVIC-181** | 2.4E+05 | 1.7E-03 | 6.9E-09 | 145.3 | 2.3 | 2.0E+05 | 1.0E-05 | 5.0E-11 | 420.0 | 7.7 | 1.1E+05 | 1.0E-05 | 8.8E-11 | 396.7 | 12.3 | 1.04E+05 | 1.00E-05 | 9.63E-11 | 448.5 | 19.1 |
| **COVIC-182** | 1.1E+05 | 5.5E-03 | 5.2E-08 | 140.1 | 3.0 | 1.8E+05 | 5.6E-05 | 3.2E-10 | 444.7 | 7.6 | 8.4E+04 | 2.2E-05 | 2.6E-10 | 463.6 | 11.4 | 9.93E+04 | 5.69E-05 | 5.73E-10 | 458.8 | 22.5 |
| **COVIC-183** | 5.0E+05 | 1.2E-02 | 2.3E-08 | 106.5 | 1.9 | 3.6E+05 | 9.4E-05 | 2.6E-10 | 365.1 | 11.4 | 1.6E+05 | 1.0E-05 | 6.3E-11 | 233.2 | 7.9 | 1.12E+05 | 9.11E-05 | 8.16E-10 | 255.5 | 10.6 |
| **COVIC-184** | 2.2E+05 | 1.8E-02 | 8.3E-08 | 140.6 | 2.4 | 2.7E+05 | 4.5E-05 | 1.7E-10 | 560.3 | 11.9 | 1.6E+05 | 7.0E-05 | 4.3E-10 | 551.2 | 17.8 | 1.81E+05 | 4.71E-04 | 2.60E-09 | 504.5 | 26.1 |
| **COVIC-185** | 3.5E+05 | 9.8E-03 | 2.8E-08 | 153.8 | 3.5 | 3.2E+05 | 3.9E-05 | 1.2E-10 | 550.6 | 14.1 | 1.8E+05 | 5.7E-05 | 3.2E-10 | 516.3 | 19.4 | 1.50E+05 | 6.26E-05 | 4.16E-10 | 536.1 | 26.1 |
| **COVIC-186** | 7.9E+04 | 1.2E-03 | 1.6E-08 | 139.0 | 1.3 | 6.8E+04 | 1.0E-05 | 1.5E-10 | 388.5 | 4.4 | 1.6E+04 | 9.1E-05 | 5.7E-09 | 481.8 | 4.2 | 8.11E+02 | 5.44E-05 | 6.70E-08 | 2739.0 | 6.9 |
| **COVIC-187** | N/A | N/A | N/A | N/A | N/A | 8.5E+04 | 1.0E-05 | 1.2E-10 | 509.2 | 3.8 | 5.8E+04 | 1.0E-05 | 1.7E-10 | 549.3 | 15.4 | 7.10E+04 | 1.00E-05 | 1.41E-10 | 580.9 | 26.0 |
| **COVIC-188** | N/A | N/A | N/A | N/A | N/A | 1.8E+05 | 2.9E-04 | 1.6E-09 | 65.9 | 2.9 | 2.5E+04 | 2.0E-04 | 7.8E-09 | 129.3 | 4.0 | 4.40E+03 | 7.82E-04 | 1.78E-07 | 735.6 | 9.4 |
| **COVIC-189** | N/A | N/A | N/A | N/A | N/A | 6.0E+04 | 1.8E-04 | 2.9E-09 | 366.7 | 4.2 | 5.9E+04 | 1.6E-04 | 2.6E-09 | 318.6 | 5.5 | 6.80E+04 | 1.00E-05 | 1.47E-10 | 274.7 | 17.9 |
| **COVIC-190** | N/A | N/A | N/A | N/A | N/A | 4.0E+04 | 1.2E-04 | 3.0E-09 | 345.1 | 4.6 | 4.2E+04 | 3.4E-04 | 8.2E-09 | 300.8 | 3.9 | 4.70E+04 | 1.89E-04 | 4.03E-09 | 262.8 | 7.8 |
| **COVIC-191** | N/A | N/A | N/A | N/A | N/A | 2.7E+05 | 9.9E-05 | 3.6E-10 | 60.0 | 2.8 | 6.5E+04 | 1.0E-05 | 1.5E-10 | 89.0 | 3.9 | 2.53E+04 | 1.00E-05 | 3.95E-10 | 176.9 | 3.6 |
| **COVIC-192** | 4.0E+05 | 9.7E-03 | 2.4E-08 | 34.8 | 1.1 | 1.5E+05 | 6.7E-05 | 4.6E-10 | 327.1 | 5.0 | 7.5E+04 | 3.1E-05 | 4.1E-10 | 380.7 | 9.9 | 9.13E+04 | 1.00E-05 | 1.10E-10 | 415.1 | 17.9 |
| **COVIC-193** | 3.0E+05 | 6.6E-03 | 2.2E-08 | 28.1 | 1.1 | 1.3E+05 | 1.9E-04 | 1.4E-09 | 268.6 | 4.1 | 7.0E+04 | 4.8E-05 | 6.9E-10 | 260.8 | 6.3 | 7.52E+04 | 1.00E-05 | 1.33E-10 | 232.3 | 25.9 |
| **COVIC-194** | 2.5E+05 | 5.5E-03 | 2.2E-08 | 52.6 | 4.2 | 1.7E+05 | 4.7E-05 | 2.7E-10 | 266.0 | 6.1 | 1.1E+05 | 3.1E-04 | 2.9E-09 | 224.3 | 6.7 | 1.05E+05 | 4.63E-04 | 4.39E-09 | 233.5 | 10.6 |
| **COVIC-195** | 2.8E+05 | 3.2E-03 | 1.1E-08 | 34.8 | 2.2 | 1.8E+05 | 1.1E-04 | 6.3E-10 | 159.7 | 2.8 | 6.5E+04 | 1.0E-05 | 1.5E-10 | 103.0 | 4.2 | 3.80E+04 | 1.00E-05 | 2.63E-10 | 113.7 | 15.0 |
| **COVIC-196** | 1.6E+05 | 1.5E-03 | 9.7E-09 | 22.7 | 2.2 | 1.8E+05 | 3.0E-04 | 1.6E-09 | 203.9 | 4.7 | 9.3E+04 | 4.3E-04 | 4.6E-09 | 227.2 | 7.0 | 1.05E+05 | 6.13E-04 | 5.81E-09 | 251.2 | 14.7 |
| **COVIC-197** | 2.8E+04 | 1.7E-02 | 5.9E-07 | 131.2 | 3.1 | 2.2E+05 | 1.2E-04 | 5.5E-10 | 322.4 | 6.4 | 1.1E+05 | 1.3E-04 | 1.2E-09 | 308.3 | 9.3 | 1.26E+05 | 2.18E-04 | 1.73E-09 | 343.0 | 16.1 |
| **COVIC-198** | 1.8E+05 | 1.3E-02 | 6.9E-08 | 64.9 | 2.0 | 2.3E+05 | 5.9E-05 | 2.5E-10 | 295.7 | 5.5 | 1.2E+05 | 5.4E-05 | 4.5E-10 | 277.9 | 9.1 | 9.95E+04 | 1.72E-04 | 1.73E-09 | 358.4 | 15.7 |
| **COVIC-199** | 1.1E+05 | 1.5E-03 | 1.5E-08 | 60.2 | 1.2 | 7.8E+04 | 1.4E-04 | 1.8E-09 | 253.0 | 2.9 | 2.2E+04 | 1.8E-04 | 8.3E-09 | 259.6 | 3.0 | 1.61E+04 | 1.00E-05 | 6.20E-10 | 216.8 | 4.3 |
| **COVIC-200** | 2.0E+05 | 9.3E-03 | 4.6E-08 | 137.5 | 3.7 | 1.7E+05 | 1.0E-04 | 6.1E-10 | 504.1 | 8.0 | 6.1E+04 | 1.0E-04 | 1.7E-09 | 483.4 | 5.7 | 1.75E+04 | 3.49E-04 | 2.00E-08 | 543.9 | 12.2 |
| **COVIC-201** | 2.0E+05 | 1.5E-03 | 7.7E-09 | 184.2 | 2.9 | 2.6E+05 | 4.1E-05 | 1.6E-10 | 501.5 | 12.6 | 1.0E+05 | 1.0E-05 | 9.6E-11 | 510.6 | 16.6 | 9.20E+04 | 2.71E-05 | 2.94E-10 | 561.5 | 22.0 |
| **COVIC-202** | 1.0E+05 | 4.1E-02 | 4.0E-07 | 313.9 | 6.9 | 3.0E+05 | 4.0E-04 | 1.3E-09 | 588.2 | 10.4 | 1.1E+05 | 7.5E-04 | 6.8E-09 | 499.7 | 18.5 | 4.05E+04 | 6.17E-04 | 1.52E-08 | 407.2 | 10.5 |
| **COVIC-REF-1** | 1.4E+05 | 3.1E-03 | 2.3E-08 | 484.9 | 4.9 | 1.7E+05 | 7.4E-05 | 4.3E-10 | 740.9 | 12.1 | 5.5E+04 | 1.0E-05 | 1.8E-10 | 742.6 | 11.9 | 1.79E+04 | 2.09E-04 | 1.17E-08 | 762.2 | 8.2 |
| **COVIC-REF-2** | 8.3E+04 | 3.1E-03 | 3.7E-08 | 424.2 | 8.2 | 1.6E+05 | 3.2E-05 | 2.0E-10 | 657.0 | 10.4 | 4.0E+04 | 1.0E-05 | 2.5E-10 | 445.7 | 12.7 | 3.56E+04 | 2.39E-05 | 6.73E-10 | 420.0 | 14.1 |

**Extended Data Table S2. Kinetic data of the High-throughput surface plasmon resonance. Related to Figure 4.**
