## Supplementary figures and images for "Structure-based design of a highly stable, covalently-linked SARS-CoV-2 spike trimer with improved structural properties and immunogenicity"

### Supplementary Figures

A

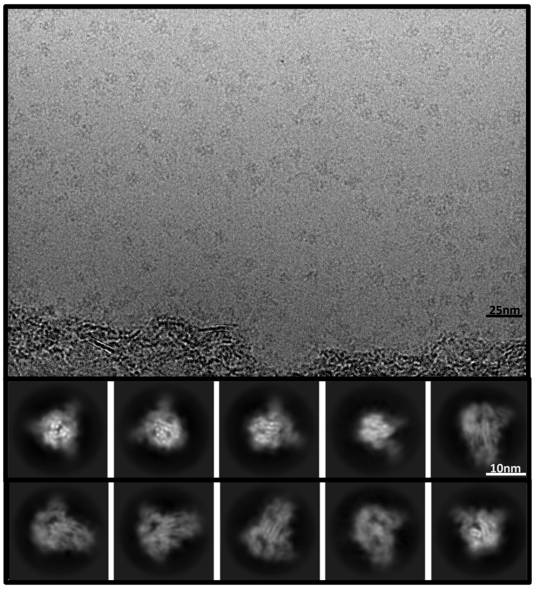


B

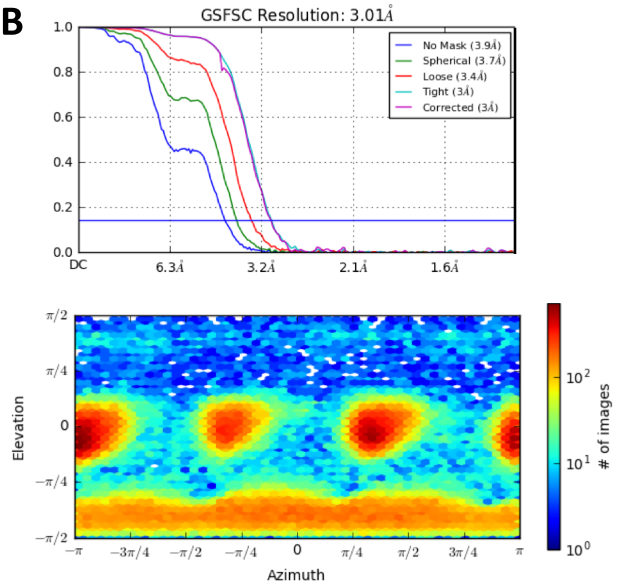


C

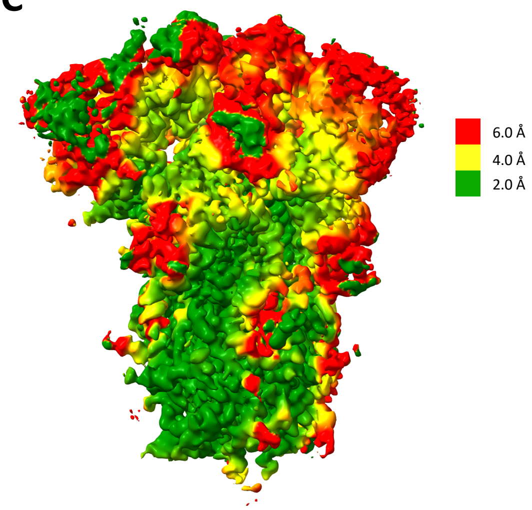


**Figure S1**

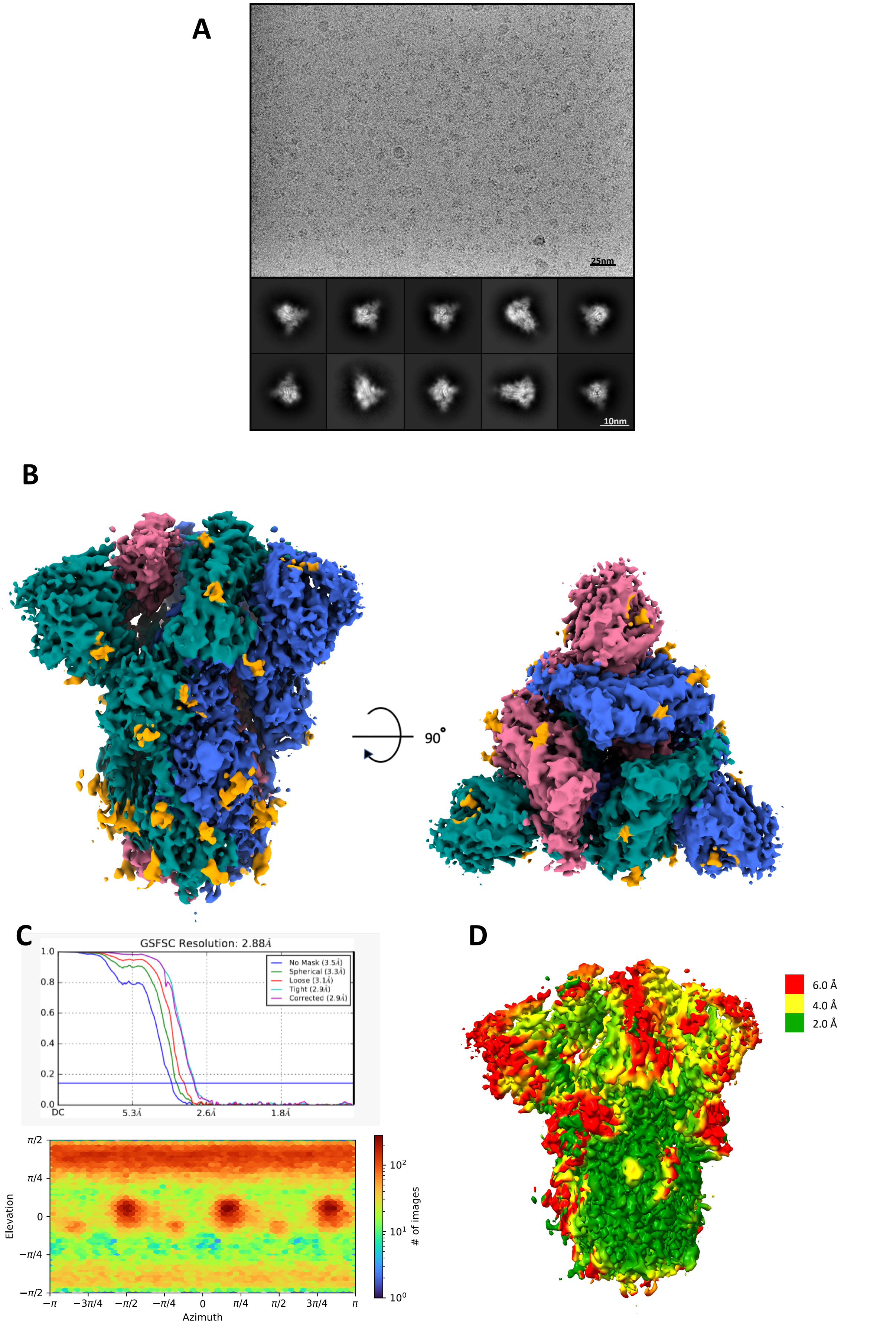

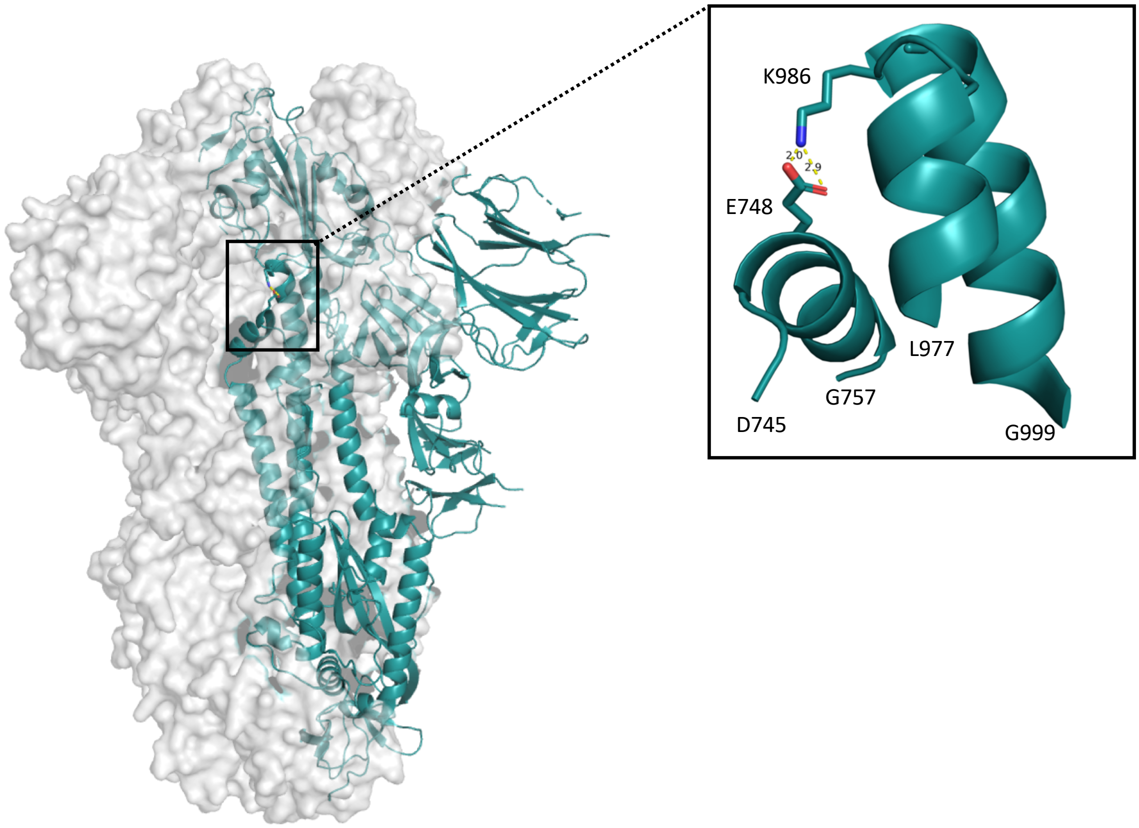


**Figure S3**

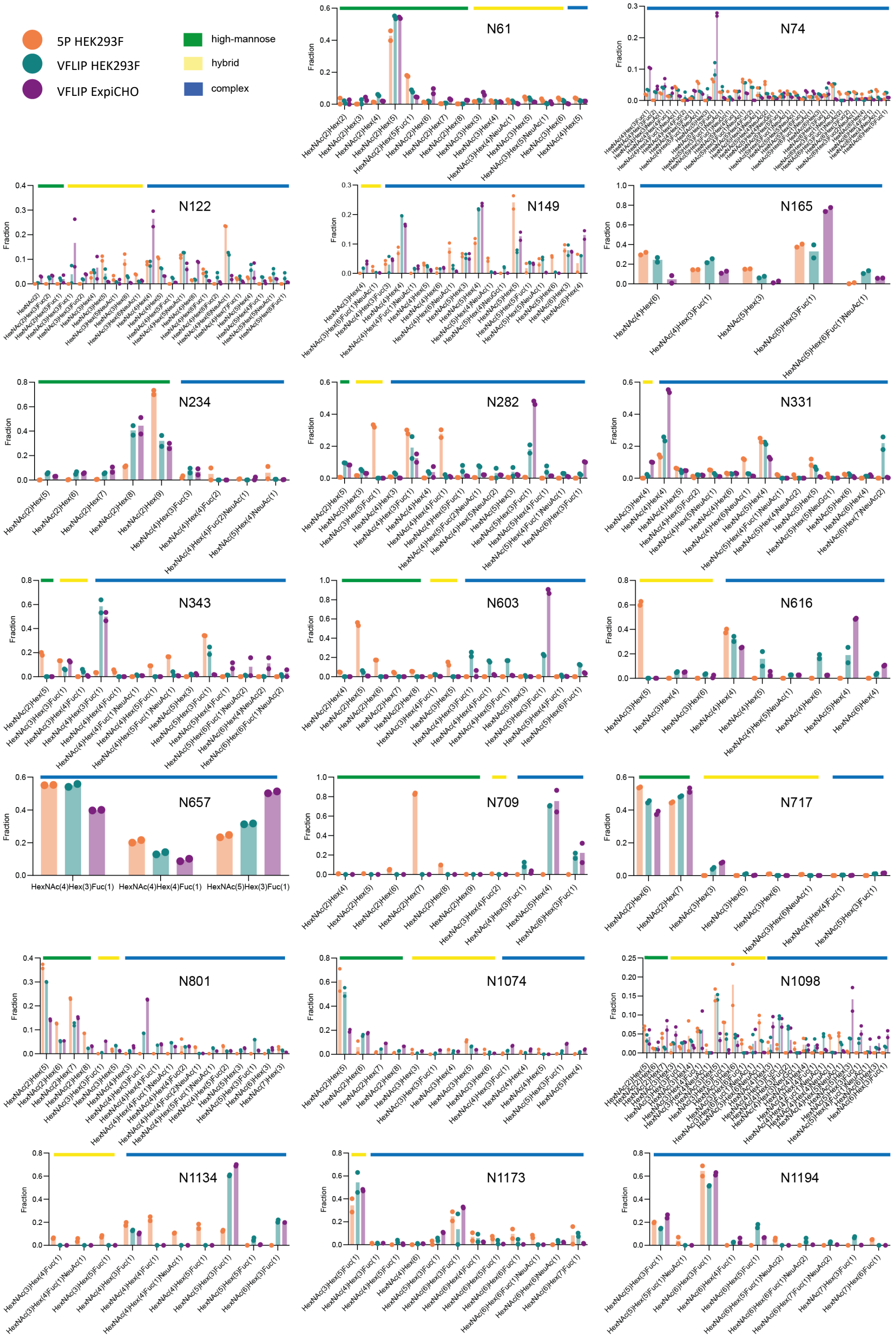

A

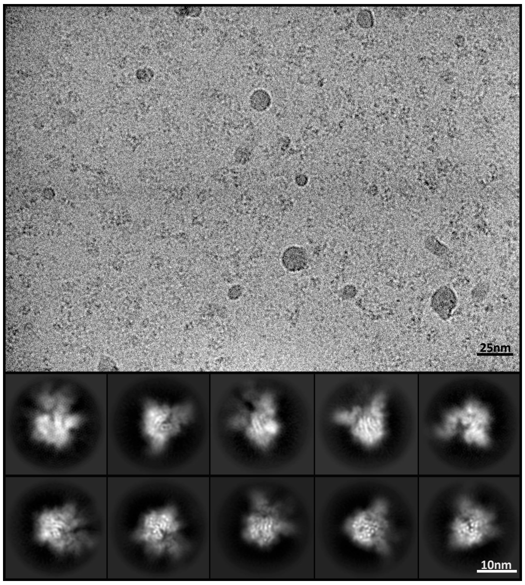


B

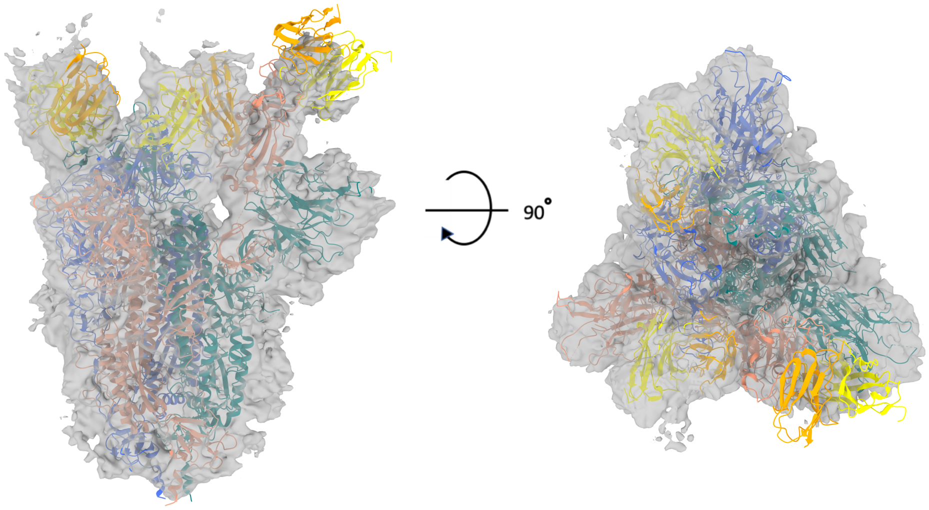


C

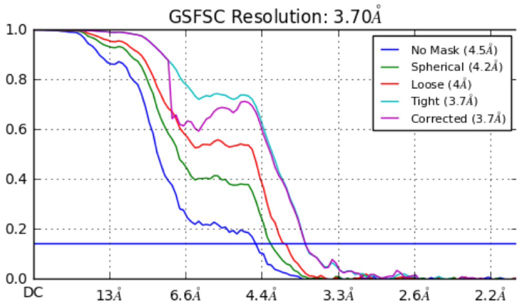


D

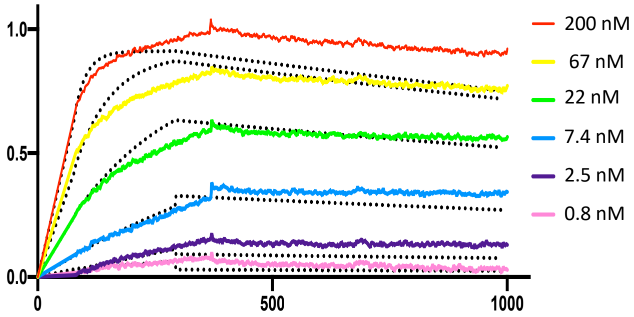


**Figure S5**

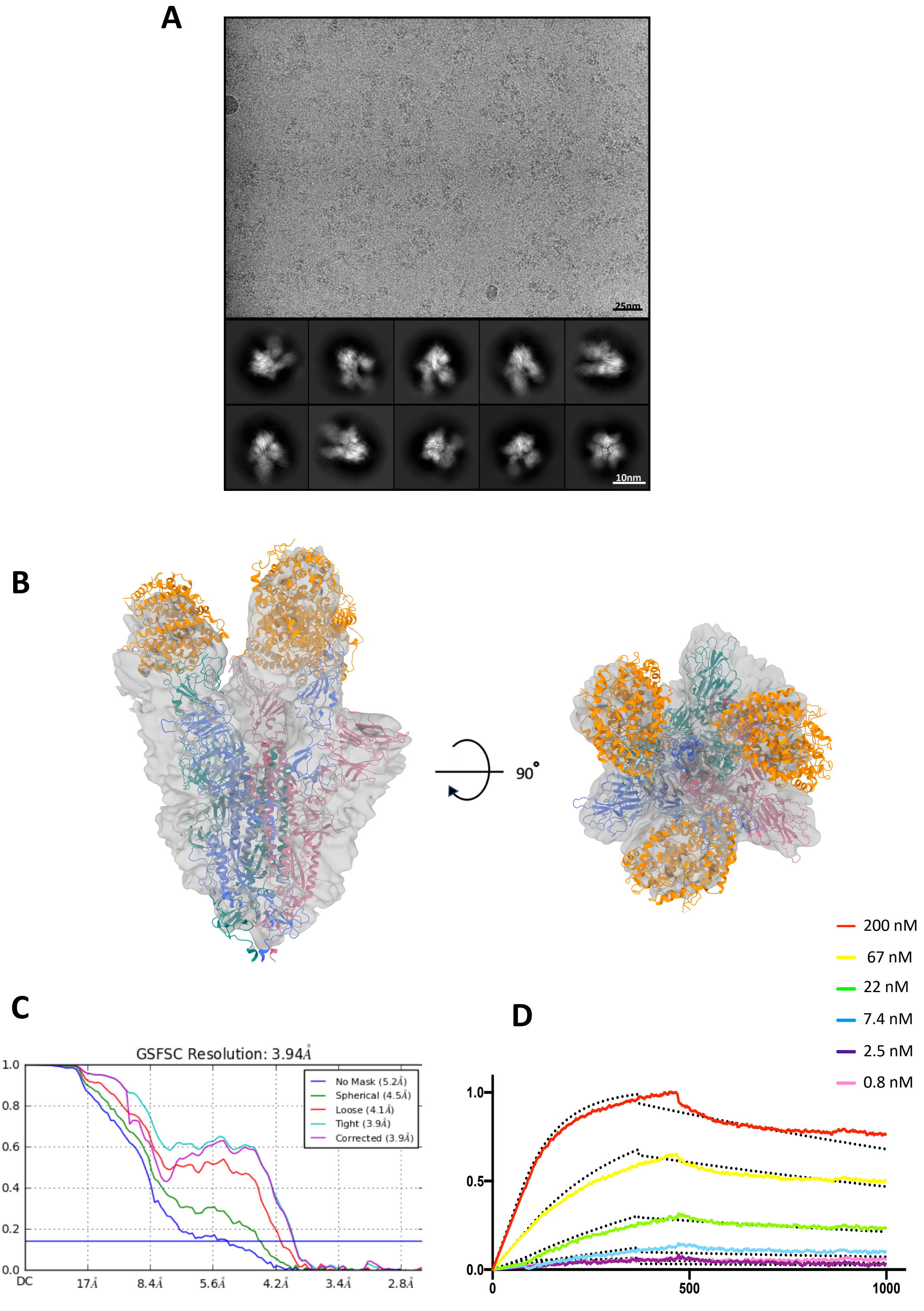

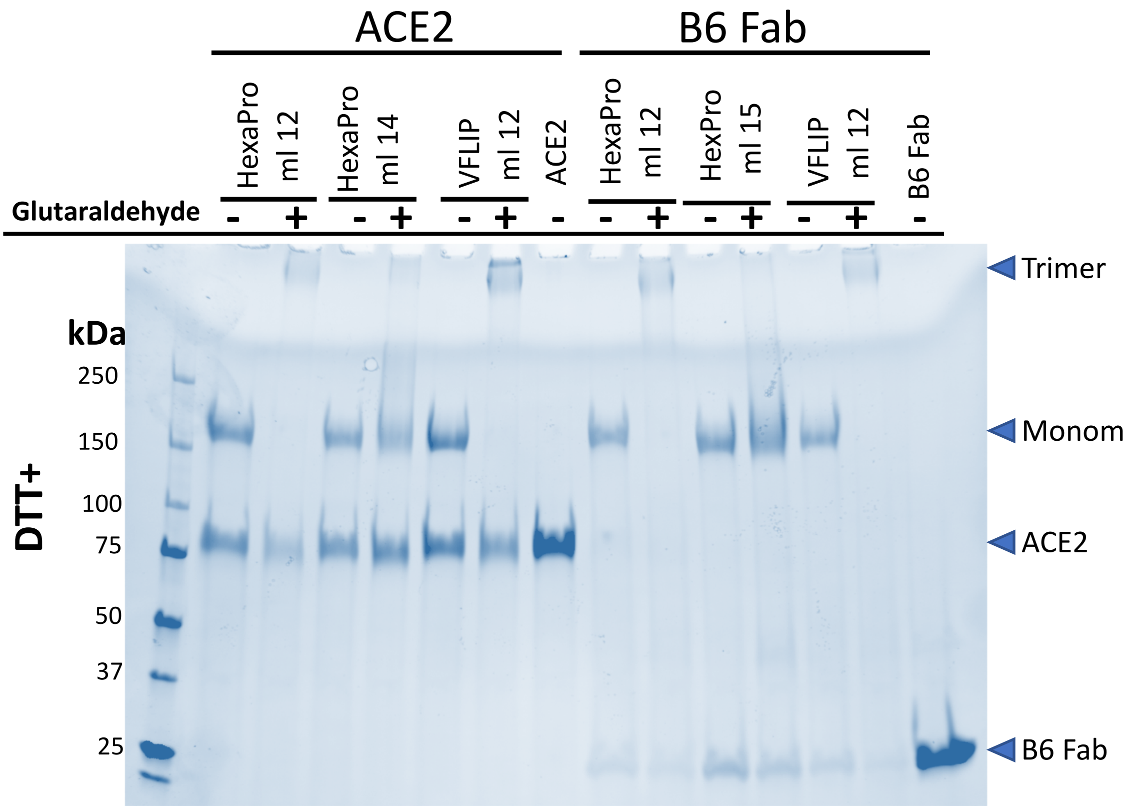

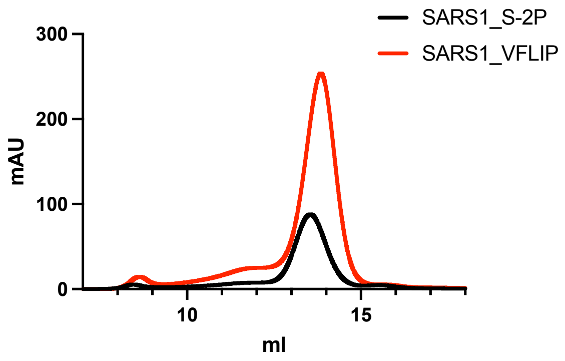

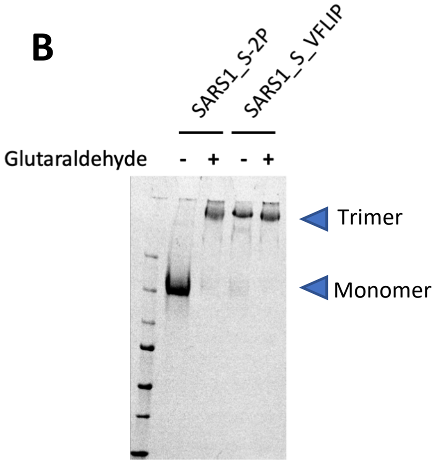
